# Homologous recombination promotes mitotic death to suppress the innate immune response

**DOI:** 10.1101/2024.05.04.592546

**Authors:** Radoslaw Szmyd, Sienna Casolin, Lucy French, Anna Gonzalez Manjon, Andrew Dhawan, Christopher B. Nelson, Léa Cavalli, Scott G. Page, Eric Hau, Hilda A. Pickett, Harriet E. Gee, Anthony J. Cesare

**Author notes:** Corresponding authors H.E.G. A.J.C.

## Abstract

Double strand breaks (DSBs) can initiate mitotic catastrophe, a complex oncosuppressive phenomenon characterized by cell death during or after cell division. Through single-cell analysis of extended live imaging, we unveiled how cell cycle-regulated DSB repair guides disparate mitotic catastrophe outcomes. Our data reveal that toxic double Holliday junctions (dHjs) generated during homologous recombination (HR) promote non-immunogenic intrinsic apoptosis in the immediate mitosis after S or G2-phase DSB induction. Conversely, the combined activity of non-homologous end joining (NHEJ), microhomology mediated end joining (MMEJ), and single strand annealing (SSA) enable G1 phase cells to tolerate high DSB loads at the cost of aberrant cell division, innate immune response activation and delayed extrinsic lethality. Targeting NHEJ, MMEJ, or SSA promotes HR-dependent mitotic death, while suppressing mitotic death fosters a robust immunogenic response. Together the data indicate that a temporal repair hierarchy, coupled with cumulative DSB load, serves as a reliable predictor of mitotic catastrophe outcomes. In this pathway, HR suppress the innate immune response by promoting mitotic lethality.

## INTRODUCTION

Mitotic catastrophe is a complex and poorly understood pathway that plays a pivotal role in oncosuppression and the clinical response to cancer therapy^1^. Within the spectrum of events that promote mitotic catastrophe are the clinical induction of genome damage including DSBs^2^. The molecular and cellular events encompassed by mitotic catastrophe include mitotic death, aberrant cell division, genomic instability, and delayed lethality resulting from mitotic errors^1,2^. Mitotic death during mitotic catastrophe is typically associated with mitotic arrest and non-immunogenic intrinsic apoptosis mediated by *BAX* and *BAK*^1, 2^. Recent evidence indicates that chromosome segregation defects following aberrant cell division activate the innate immune response through cGAS DNA sensing and/or MAVS RNA sensing pathways^3–6^. This can potentiate Interferon regulatory factor 3 (IRF3) and STAT1-mediated interferon stimulated gene (ISG) expression, paracrine signaling, and extrinsic lethality^3, 6^. Despite the common use of the term mitotic catastrophe to describe various mitosis-centric outcomes following genomic damage, the molecular mechanisms guiding the diverse outcomes within this phenomenon remain unknown. Specifically, it is unclear what distinguishes cells that die in mitosis from those that survive cell division. Given that this cellular decision may underly the induction of non-immunogenic versus immunogenic outcomes, we consider it a focus of central therapeutic importance.

DSBs are repaired through four cycle-dependent pathways^7, 8^. NHEJ operates throughout the cell cycle, while HR and SSA are active in S and G2^9, 10^. MMEJ is active outside of G1^11, 12^. NHEJ is generally considered to be the only DSB repair pathway active in G1; however, recent evidence suggests limited G1 phase HR^13^. NHEJ operates independently of DNA end resection, as DNA-dependent protein kinase catalytic subunit (DNA-PKcs) and Ligase 4 (LIG4) cooperate to covalently ligate DNA ends^14^. In contrast, the other DSB repair mechanisms require DNA end-resection to expose single-strand (ss) DNA sequences that are used in the repair process^15^. During MMEJ and SSA, homologous ss sequences align, followed by removal of the remaining non-homologous sequence, polymerase-mediated fill-in, and ligation^10, 16^. SSA relies on RAD52 while Polθ is indispensable for MMEJ. Both MMEJ and SSA are error-prone, often leading to chromosomal translocations and/or large-scale deletions^10, 16^.

HR uses complementary sequence from the sister chromatid, or less frequently, the homologous chromosome, as templates for error-free repair. Biochemically, HR represents the most intricate DSB repair pathway^17, 18^. In the HR process, the DNA damage response kinase Ataxia telangiectasia and Rad3 related (ATR) phosphorylates PALB2, promoting its interaction with BRCA2 and facilitating the recruitment of BRCA2-PALB2 to DSBs^19, 20^. Subsequently, BRCA2 loads RAD51 onto resected ssDNA, leading to the formation RAD51-coated ssDNA filaments^21, 22^ that promote strand-invasion and establish a displacement-loop (D-loop) within homologous DNA sequence^23^. Following D-loop formation, HR can proceed through non-crossover synthesis-dependent strand annealing (SDSA) reactions or progress to form dHJs that may promote crossover repair^24^. RTEL1 suppresses dHJ formation to promote SDSA^25^, and dHJs are resolved through several different mechanisms including SLX4 mediated endonucleolytic cleavage^26–28^. Due to its error-free nature, HR is often regarded as the preferred DSB repair mechanism.

In this study, we utilized long-duration live imaging and single-cell analysis of human cells expressing the three-color FUCCI (3F) cell cycle reporter to investigate mitotic catastrophe in response to genome damage. We have identified that different mitotic catastrophe outcomes in p53 compromised cells are regulated by the choice of DSB repair pathways. The formation of toxic dHJs emerges as the key determinant that dictates whether a cell perishes through immunogenic or non-immunogenic pathways.

## RESULTS

### Genomic damage promotes distinct cell cycle-regulated mitotic catastrophe outcomes

To visualize all mitotic catastrophe outcomes following genome damage with single cell resolution, we used live fluorescent imaging of p53 compromised HeLa cervical carcinoma cells expressing the 3F cell cycle reporter (Fig. 1A, B and Supplementary Video 1)^29^. We treated cultures with single fractions of 2 to 20 Gy ionizing radiation (IR) and recorded outcomes through images captured at six-minute intervals for up to 120 hours. We noted cell cycle phase at the time of mock or IR treatment, and broadly classified cell cycle outcomes as cell division, interphase death, or mitotic death (Fig. 1B and Supplementary Video 2). Cell division included normal mitosis and completed aberrant mitoses containing chromosome segregation errors, micronuclei, multipolar spindles, multipolar divisions, cytokinesis defects, and/or mitotic slippage (mitosis to interphase transition without anaphase or telophase) (Extended Data Fig. 1A and Supplementary Video 2).

**Figure 1:**
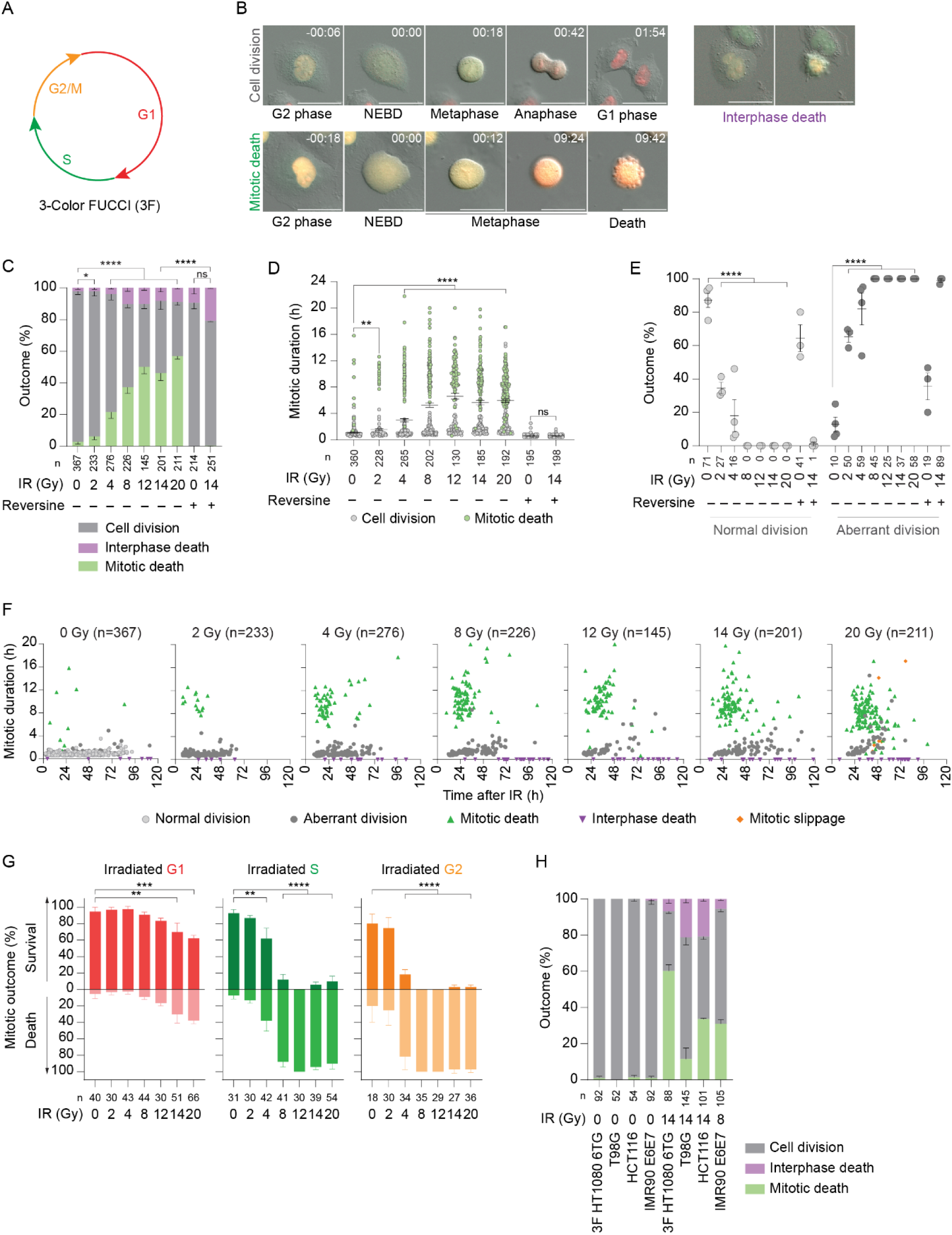
Distinct waves of cell death occur during mitotic catastrophe as a function of cell cycle phasing at damage induction. **A)** Schematic of 3-color FUCCI coloration ^29^. **B)** Representative images from live microscopy of 3F HeLa cultures. Time is hrs:min relative to nuclear envelope breakdown (NEBD). Scale bar = 50µm. **C, D)** Outcome (C) (mean ± SEM of ≥ 3 biological replicates; Fisher’s exact test; p values for mitotic death are shown) and mitotic duration (D) (mean ± SEM of ≥ 3 biological replicates; Kruskal Wallis multiple comparison test) of 3F HeLa cells measured through 120 hours of live imaging immediately following mock or IR treatment. Cells are color coded for mitotic survival or death in (D). **E)** Outcomes of completed mitoses from 3F HeLa cultures during the first cell division after mock or IR treatment (mean ± SEM of ≥ 3 biological replicates; Fisher’s exact test). **F)** Multi-dimensional representation of the data from (C). Each symbol represents an individual event. Symbol position relative to the x-axis is time after mock or IR treatment that mitotic entry or interphase death occurred. Y-axis indicates mitotic duration to cell division or death. **G)** First mitosis outcome measured by live imaging of 3F HeLa cultures following mock or IR treatment as a function of cell cycle phase at irradiation (mean ± SEM of ≥ 3 biological replicates; Fisher’s exact test). **H)** Outcome as in (C) of multiple cell lines following mock or IR treatment (mean ± SEM of = 2 biological replicates). Where appropriate, ns = not significant, *p < 0.05, **p < 0.01, *** p <0.001, ****p < 0.0001. For panels C – H, n = number of cells quantified across all biological replicates. For panels C – E, where indicated 0.5µM Reversine was added two hours prior to mock or 14 Gy IR treatment.

We observed IR dose-dependent increases in, and a strong correlation between, mitotic duration and mitotic death (Fig. 1C, D and Extended Data Fig. 1B). We note that even with a low dose of 2 Gy, mitotic death was significantly increased (Extended Data Fig 1B). Higher dosages of 8 or more Gy conferred ubiquitous aberrant cell division and similar levels of interphase lethality during the 120-hour observation window (Fig. 1C, E). Single-cell outcomes were contextualized in multiple dimensions by comparing time after IR, mitotic duration, and fate (Fig. 1F). This demonstrated distinct grouping of cell death outcomes. Specifically, cells that arrested and died in the first mitotic attempt 24 to 48 hours post-IR and cells that died predominantly in interphase 40 hours or more post-IR. Interphase death occurred ≥ 86 ± 9% (mean ± SEM) of the time following at least one aberrant division (Extended Data Fig. 1C). MPS1 kinase regulates the Spindle Assembly Checkpoint (SAC) which prevents mitotic exit until chromosomes correctly align and establish tension across the mitotic spindle^30^. The MPS1 inhibitor Reversine^31^ prevented mitotic arrest and mitotic death in 14 Gy irradiated 3F HeLa cultures (Fig. 1C – E and Extended Data Fig. 1B) confirming that mitotic lethality following lethal genomic damage was dependent upon persistent SAC activation.

3F HeLa cultures revealed a strong correlation between cell cycle phase at irradiation and mitotic catastrophe outcomes (Fig. 1A, G). Treatment of asynchronous cultures with 4 Gy caused significant mitotic death in cells irradiated during S and G2-phase (Fig. 1G). With 8 or more Gy, S and G2 irradiated cells died during the first attempted cell division in 88 ± 6% of occurrences (Fig. 1G). This was surprising as mitotic death in solid tumors was previously considered a rare outcome^32^. Conversely, even with 20 Gy, cells irradiated in G1 predominantly survived their current cell cycle and completed mitosis, only to succumb thereafter, primarily in interphase (Extended Data Fig. 1D).

We performed similar live imaging in a panel of cell types; primary IMR90 fibroblasts, IMR90 fibroblasts expressing HPV16 E6 and E7 (IMR90 E6E7) to respectively compromise p53 and Rb ^33^, p53 mutant HT1080 6TG fibrosarcoma and T98G glioblastoma cells, p53 knockout HCT116 colorectal carcinoma cells, and p53 wild type (WT) A549 lung adenocarcinoma cells (Fig. 1H and Extended Data Fig. 1E-G). The A549 and HT1080 6TG cultures expressed the 3F cell cycle reporter, and IMR90 fibroblasts the first generation 2-color FUCCI (2F) system^34^ (Extended Data Fig. 1G). All irradiated p53-compromised cultures exhibited a response consistent with HeLa (Fig. 1H). Specifically, a grouping of cells that arrested and died in the first mitosis (20 – 65% of outcomes; mean ± SEM: 44 ± 9%), and a cell grouping that survived the first mitosis and perished thereafter, mostly in interphase (12 – 63% of outcomes; mean ± SEM: 38 ± 11%). 3F HT1080 6TG cultures also displayed the same strong correlation as 3F HeLa between cell cycle phase at irradiation and first mitosis outcome (Extended Data Fig. 1E). In contrast, similar IR doses induced a sustained G0/G1 arrest in p53 WT 2F IMR90 and 3F A549 cells (Extended Data Fig. 1F, G). For p53 WT cells in and/or entering S/G2 following IR, G0/G1 arrest was preceded by mitotic bypass (100% of IMR90 and 52% of A549 occurrences) or a completed cell division (48% of A549 occurrences) (Extended Data Fig. 1F, G).

Consistent with previous descriptions of mitotic catastrophe, we observed several distinct outcomes following lethal IR. Live 3F imaging, however, revealed the previously unrecognized quality that distinct cell death outcomes were a function of cell cycle phase at irradiation. We term mitotic death in the first attempted cell division immediate mitotic death, and cell death following at least one cell division delayed lethality.

### DSB repair regulates how cells die during mitotic catastrophe

Given the strong relationship between cell cycle phase at irradiation and the mode of cell death, we hypothesized that mitotic catastrophe outcomes were DSB repair related (Extended Data Fig. 2A). To suppress NHEJ we targeted DNA-PKcs with 0.5µM of the specific inhibitor NU7441^35^, and validated with the CRISPR-mediated repair assay^36^ (Extended Data Fig. 2B and Supplementary Video 3). NU7441 did not affect non-irradiated cultures, nor the survival of S and G2 irradiated cells (Fig. 2A, B). NU7441 did, however, significantly induce immediate mitotic death in G1 cells treated with 20 Gy IR (Fig. 2A, B and Extended Data Fig. 2C). Genetic depletion of DNA-PKcs or LIG4 conferred similar levels of immediate mitotic death in 14 Gy irradiated G1 cells (Extended Data Fig. 2D-F).

**Figure 2:**
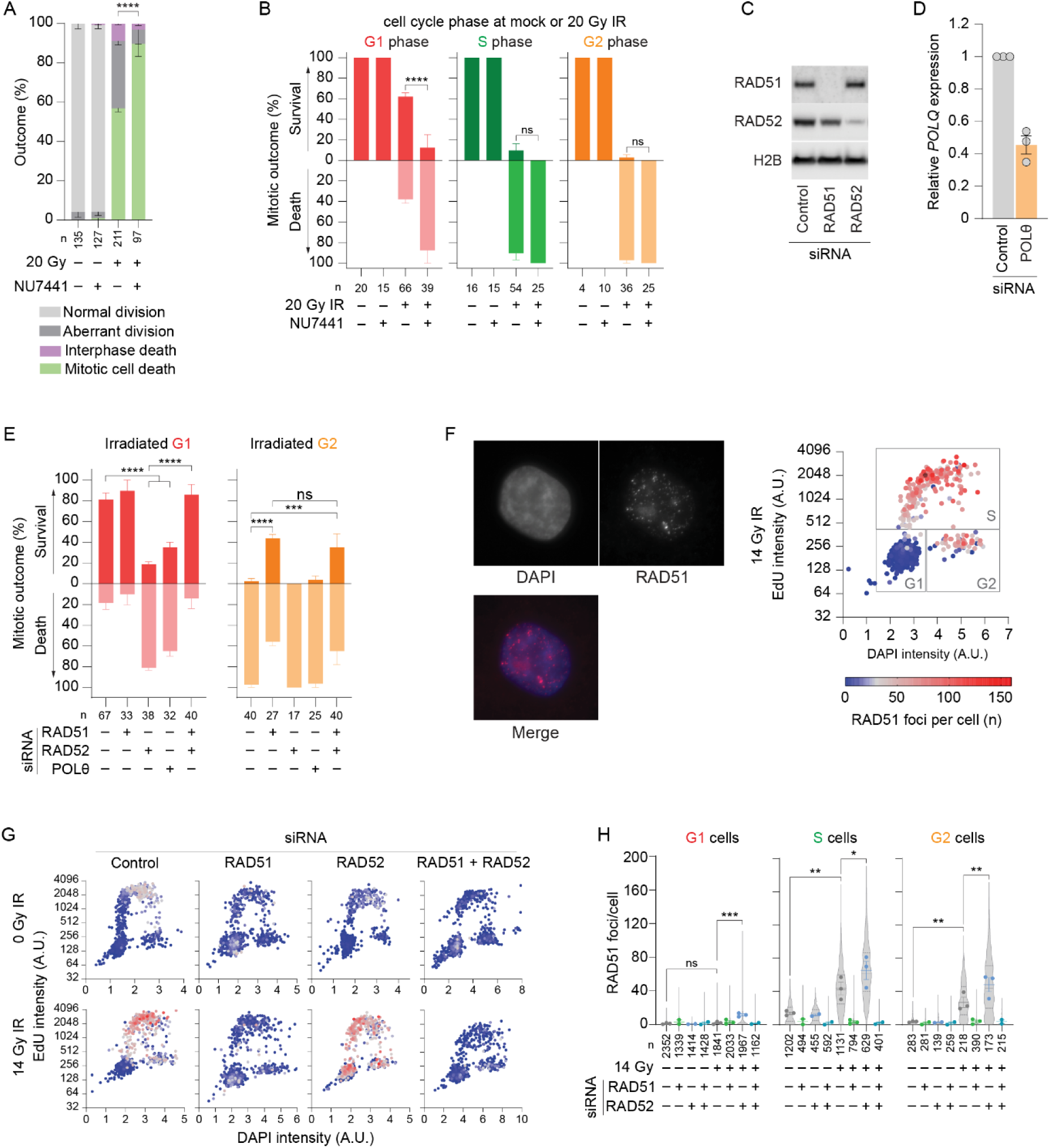
Double strand break repair pathways regulate how cells perish during mitotic catastrophe. **A)** Outcome measured by live imaging of 3F HeLa cultures following mock or 20 Gy IR treatment ± 0.5µM NU7441 added 30 minutes prior to treatment (mean ± SEM of ≥ 2 biological replicates; Fisher’s exact test; p values for changes in mitotic death are depicted). The 20 Gy IR sample without NU7441 are the same data as in Fig. 1C. **B)** Data from (A) classified by first mitosis outcome as a function of cell cycle phase at irradiation (mean ± SEM of ≥ 2 biological replicates; Fisher’s exact test). **C)** Western blots of whole cell extracts derived from 3F HeLa cultures treated with the indicated siRNAs. **D)** RT-qPCR measurement of *POLθ* mRNA from siRNA transfected 3F HeLa cells (mean ± SEM; n = 3 biological replicates). Fold change in gene expression is relative to control siRNA and normalized to 1. **E)** First mitosis outcome measured as in (B) in 3F HeLa cells transfected with the indicated siRNAs and irradiated with 14 Gy IR (mean ± SEM of ≥ 3 biological replicates; Fisher’s exact test). **F)** Representative images (left) and quantitation of RAD51 immunofluorescence in HeLa cells as a function of the cell cycle (right) using multidimensional fluorescent microscopy. Each dot represents a single cell. G1, S, and G2 classifications are based on EdU and DAPI intensity. **G)** RAD51 foci in HeLa cells transfected with the indicated siRNA ± 14 Gy IR, imaged and presented as in (F). A representative biological replicate is shown. **H)** Quantitation of the experiment in (G) (mean ± SEM of ≥ 2 biological replicates; ordinary one-way ANOVA with Fisher’s LSD test; violin plots depict all datapoints per condition, replicate means are shown by individual points, dotted lines mark quartiles, and solid lines represent median values). Where appropriate, ns = not significant, *p < 0.05, **p < 0.01, *** p <0.001, ****p < 0.0001. For panels A, B, E, and H, n = number of cells quantified across all biological replicates.

We next depleted RAD51, RAD52, and/or Polθ, factors respectively essential for HR, SSA, and MMEJ^37, 38^ (Fig. 2C, D and Extended Data Fig. 2G). SiRNA effects were validated using I-SceI HR or SSA reporter constructs^38, 39^ and/or the CRISPR-mediated repair assay^36^ (Extended Data Fig. 2B, H). RAD51 knockdown rescued immediate mitotic death in G2 cells irradiated with 14 Gy (Fig. 2E, Extended Data Fig 2I, and Supplementary Video 4), implicating HR in mitotic lethality. Neither RAD52 nor Polθ depletion influenced S/G2 irradiated cell outcomes (Fig. 2E and Extended Data Fig. 2I). Because HR, SSA, and MMEJ activity are limited to S and G2, we predicted that targeting these pathways would not affect G1 irradiated cells. Surprisingly, RAD52 or Polθ depletion resulted in a significant induction of immediate mitotic death in 14 Gy irradiated G1 cells (Fig. 2E). RAD51 depletion alone had no effect on G1 following IR (Fig. 2E). However, relative to singular RAD52 siRNA, co-depleting RAD51 and RAD52 rescued immediate mitotic lethality in G1, S and G2 irradiated populations, again implicating HR in mitotic death (Fig. 2E and Extended Data Fig. 2G). Co-depleting RAD51 and Polθ, or triple RAD51, RAD52, and Polθ knockdown failed to rescue mitotic death compared to Polθ siRNA alone (Extended Data Fig. 2G, I), potentially reflecting the RAD51 and Polθ synthetic lethal relationship^40^.

Previous efforts demonstrated competition between RAD51 and RAD52^8, 10^ and indicated IR doses above 4 Gy induced a switch from error-free RAD51 gene conversion to mutagenic RAD52-dependent SSA^41^. We measured RAD51 immunofluorescence within the cell cycle context (Fig. 2F). RAD51 foci were present in non-irradiated S phase cells consistent with spontaneous replication stress^42, 43^. 14 Gy IR induced RAD51 foci in S and G2 phase consistent with cell cycle regulated HR, and RAD51 siRNA suppressed RAD51 foci across all conditions (Fig. 2G, H). When combined with RAD52 siRNA, 14 Gy IR conferred elevated RAD51 foci in S and G2, and unexpectedly induced G1 phase RAD51 foci (Fig. 2G, H). This is consistent with RAD51 and RAD52 competing for repair and indicates that HR is prematurely engaged in irradiated RAD52 depleted cultures that experience a significant increase in RAD51-dependendent immediate mitotic death (Fig. 2E). RAD51 therefore promotes immediate mitotic death during mitotic catastrophe. In contrast, DNA-PKcs, LIG4, Polθ, and RAD52 cooperate to confer first cell cycle survival in G1 irradiated cells and facilitate subsequent delayed lethality.

### Double Holliday junctions promote mitotic death

To further explore HR in in the context of mitotic catastrophe, we examined additional recombination components. Consistent with their role promoting RAD51 filaments, BRCA2 or PALB2 depletion prevented RAD51 foci formation in 14 Gy irradiated cells (Fig. 3A and Extended Data Fig. 3A). We enriched G2 cells by synchronizing RAD51-, PALB2-, or BRCA2-depleted 3F HeLa cultures at the G1/S boundary using a double thymidine block, released, irradiated six hours later, and visualized outcomes through live imaging. BRCA2, PALB2, and RAD51 siRNA all significantly rescued mitotic arrest and immediate mitotic death in G2 irradiated cells (Fig. 3B, C).

**Figure 3:**
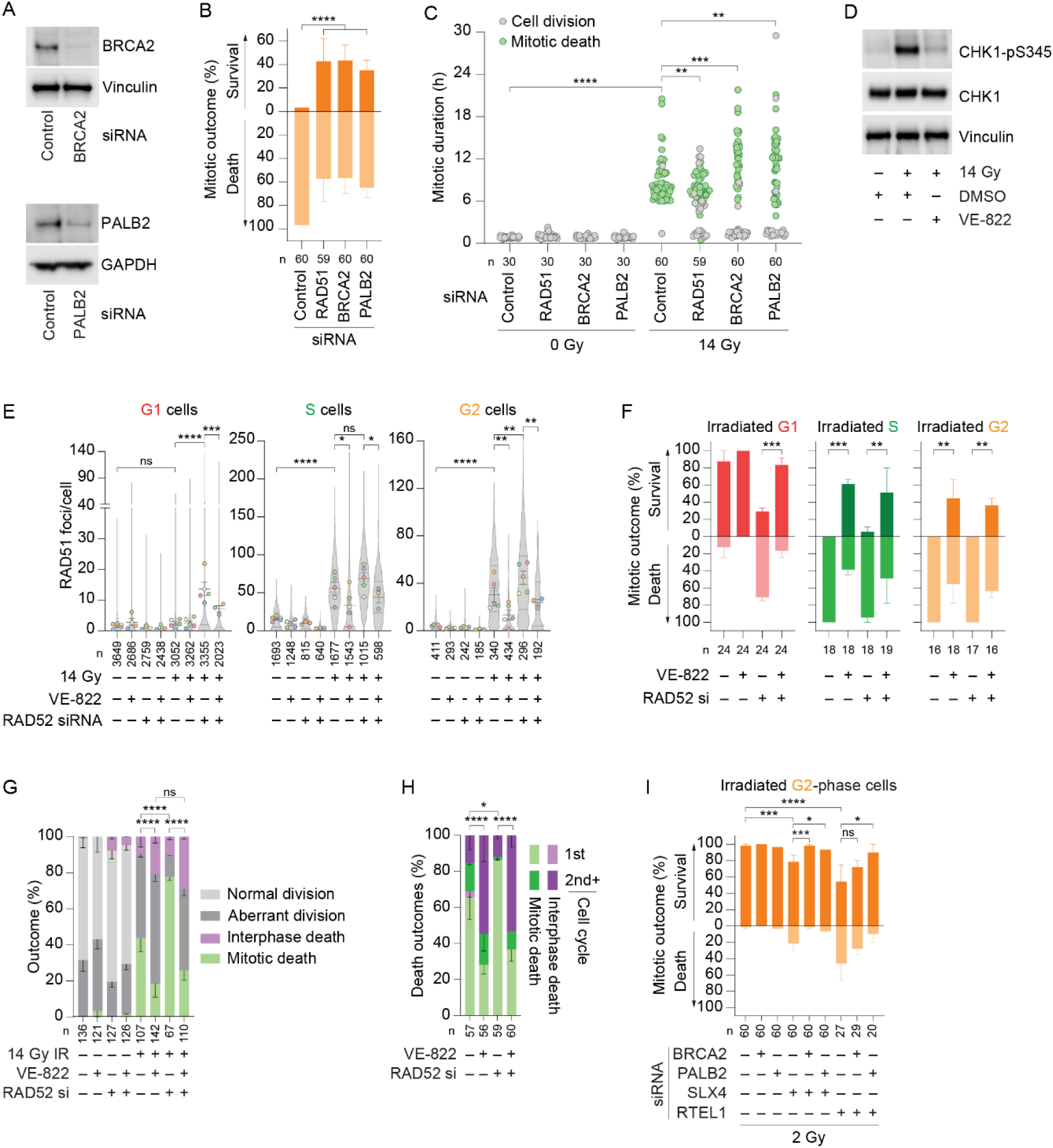
Toxic double Holliday junctions promote immediate mitotic death. **A)** Western blots of whole cell extracts derived from 3F HeLa cells treated with the indicated siRNAs. **B)** First mitosis outcome measured by live imaging following 14 Gy IR of synchronized G2-irradiated 3F HeLa cells transfected with the indicated siRNAs (mean ± SEM of 2 biological replicates; Fisher’s exact test). **C)** Mitotic duration of 3F HeLa cells treated with the indicated siRNAs following mock or 14 Gy IR (all individual datapoints from 2 biological replicates are shown; Kolmogorov-Smirnov test). **D)** Western blots of whole cell extracts derived from HeLa cells one hour following mock or 14 Gy IR ± 0.2µM VE-822. **E)** Quantification of RAD51 foci in HeLa cells measured through multidimensional fluorescent microscopy as shown in Fig. 2F from siRNA transfected cultures ± 0.2µM VE-822 (mean ± SEM of ≥ 3 biological replicates; ordinary one-way ANOVA with Fisher’s LSD test; data are presented as in Fig. 2H). **F)** First mitosis outcome in asynchronous siRNA transfected 3F HeLa cells measured by live imaging as a function of cell cycle phase at irradiation for cultures treated with 14 Gy IR ± 0.2µM VE-822 (mean ± SEM of 2 biological replicates; Fisher’s exact test). **G)** Cellular outcomes from the experiment in (F) throughout the 120-hour imaging duration (mean ± SEM of 2 biological replicates; Fisher’s exact test; p values for mitotic death are depicted). **H)** Cell death outcomes in 14 Gy irradiated samples from (G) shown as a percentage of all death events (mean ± SEM of 2 biological replicates; Fisher’s exact test; p values are for an overall death in the 1st vs 2nd+ cell cycle). **I)** First mitosis outcome measured by live imaging following 2 Gy IR of synchronized G2-irradiated 3F HeLa cells transfected with the indicated siRNAs (mean ± SEM of 2 biological replicates; Fisher’s exact test). For D – H, VE-822 was added 1 hour prior to mock or IR treatment. Where appropriate, ns = not significant, *p < 0.05, **p < 0.01, *** p <0.001, ****p < 0.0001. For panels B, C, and E-H, n = number of cells quantified across all biological replicates.

ATR regulates diverse genome stability-related activities including HR^19^. We suppressed ATR activity with the specific inhibitor VE-822^44^, as indicated by diminished phosphorylation of the downstream effector CHK1 (Fig. 3D). I-SceI DSB repair reporter constructs showed that VE-822 strongly suppressed HR while only mildly inhibiting SSA (Extended Data Fig. 3B, C). We treated 3F HeLa cultures with RAD52 siRNA to increase HR activity, in the presence or absence of VE-822. Congruent with ATR regulating mitotic lethality via HR, VE-822 suppressed RAD51 foci and immediate mitotic lethality in S and G2 irradiated cells from control siRNA samples, and in G1, S and G2 irradiated cells within RAD52 depleted cultures (Fig. 3E, F and Extended Data Fig. 3D). Under these conditions, mitotic death rescue was accompanied by a corresponding increase in delayed interphase lethality (Fig. 3G, H). The data indicate that when RAD52 is present, ATR-dependent HR promotes mitotic death in irradiated S and G2 cells, and that this outcome extends to G1-irradidated cells when RAD52 is depleted.

Downstream of RAD51-mediated strand invasion, HR proceeds through either SDSA or dHJ formation^24^. RTEL1 promotes SDSA by suppressing second end capture and dHJ formation^25^, while SLX4 supports dHJs resolution via endonucleolytic cleavage^26–28^. We depleted RTEL1 or SLX4 alone or in combination with PALB2 or BRCA2 in 3F HeLa cells (Extended Data Fig 3E, F), enriched for G2 as described above, and irradiated with a low dose of 2 Gy. Both RTEL1 and SLX4 siRNA conferred a significant increase in immediate mitotic death in G2 irradiated cells that was rescued when HR initiation was suppressed by BRCA2 and/or PALB2 co-depletion (Fig. 3I and Extended Data Fig. 3G, H). The data indicate that mitotic arrest and death during mitotic catastrophe results from unresolved dHJs. We also note that mitotic death becomes more prevalent with low dose IR under genetic conditions where dHJs persist.

### Chromosome structural rearrangements correlate with mitosis survival

Cytogenetic preparations from 14 Gy irradiated HeLa cultures revealed chromosome structural aberrations including S/G2 repair-mediated radial chromosomes and chromatid-type fusions (Fig. 4A and Extended Data Fig. 4A), and G1 NHEJ mediated multi-centric chromosome-type fusions and circular chromosomes (Extended Data Fig. 4B)^45^. Radials and chromatid-type fusions were prevalent at 24- and 36-hours post-IR (Fig. 4B and Extended Data Fig. 4C), contemporaneous with the first mitosis after irradiation (Fig. 1F). RAD51 depletion, alone or in combination with RAD52 siRNA, rescued mitotic death following 14 Gy IR (Fig. 2E), but increased mitotic cells containing radials or chromatid-type fusions (Fig. 4C and Extended Data Fig. 4D). Conversely, RAD52 or Polθ depletion suppressed chromatid-type fusions and radial chromosomes (Fig. 4C and Extended Data Fig. 4D) but induced mitotic lethality in the G1 irradiated population (Fig. 2E). NHEJ, MMEJ, and SSA-dependent chromosomal aberrations therefore correlate with mitotic survival.

**Figure 4:**
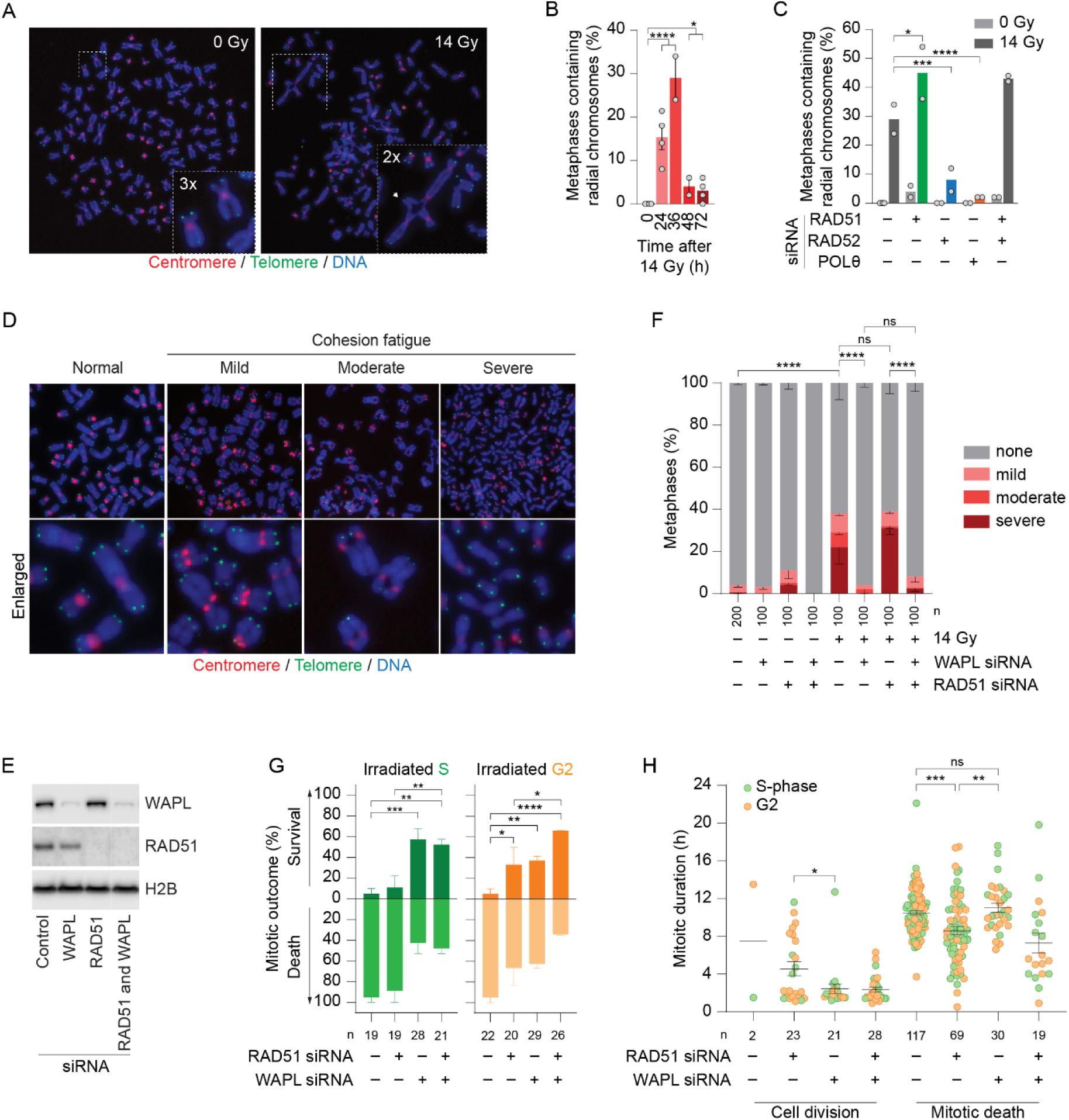
Chromosomal structural aberrations correlate with first mitotic survival during mitotic catastrophe while cohesion fatigue promotes mitotic death. **A)** Representative images of cytogenetic chromosome spreads from HeLa cells stained with DAPI (blue), and centromere (red) and telomere (green) fluorescent in situ hybridization (FISH). A radial chromosome is shown on the right. **B)** Quantitation of radial chromosomes from control siRNA treated HeLa cultures at the indicated times following 14 Gy IR (mean ± SEM of ≥ 2 biological replicates scoring 50 metaphases per replicate; Fisher’s exact test). **C)** Quantification of radial chromosomes in HeLa culturtreated with the indicated siRNAs ± 14 Gy IR (mean ± SEM; n = 2 biological replicates; Fisher’s exact test). The control siRNA data are the 0- and 36-hour time points from (B). **D)** Representative images of cohesion fatigue in HeLa cytogenetic preparations prepared as in (A). **E)** Western blots of whole cell extracts derived from siRNA transfected HeLa cells. **F)** Quantitation of cohesion fatigue in siRNA transfected HeLa cultures ± 14 Gy IR (mean ± SEM of ≥ 2 biological replicates scoring 50 metaphases per condition; Fisher’s exact test). **G)** Outcome of the first mitosis in 14 Gy irradiated S and G2 cells within asynchronous 3F HeLa cultures as determined by live imaging (mean ± SEM of ≥ 2 biological replicates; Fisher’s exact test). **H)** Mitotic duration of cells from (G) color coded by cell cycle phase at irradiation (n ≥ 2 biological replicates; Kolmogorov-Smirnov test). Where appropriate, ns = not significant, *p < 0.05, **p < 0.01, *** p <0.001, ****p < 0.0001. For panels F – H, n = number of cells quantified across all biological replicates.

### Cohesion fatigue promotes mitotic death

Cytogenetic preparations from irradiated cultures also revealed evidence of cohesion fatigue^46^ (Fig. 4D, E). Cohesion fatigue results from microtubule pulling forces exerted during mitotic arrest that render asynchronous sister chromatid separation, initiating at the centromeres and moving distally telomeric, mediated by the cohesion antagonist Wings apart like protein homologous (WAPL)^47^. We classified mild cohesion fatigue as visible separation of sister-centromeres, moderate as separation of centromere adjacent regions while telomeres remain cohesive, and severe as complete chromatid separation (Fig. 4D)^47^. Cohesion fatigue progression from mild to severe is a function of mitotic arrest duration^47^.

Depleting WAPL in 14 Gy irradiated 3F HeLa cultures suppressed cohesion fatigue and rescued immediate mitotic death in S and G2 irradiated cells (Fig. 4E-G and Extended Data Fig. 4E). WAPL was recently implicated in enabling RAD51 access to the chromatin during replication stress repair^48^, suggesting WAPL may enable HR to promote mitotic death. We observed a subtle reduction in HR with WAPL depletion (Extended Data Fig. 4F). However, the distinct phenotypes accompanying individual RAD51 or WAPL depletion suggested independent mechanisms of mitotic death rescue (Fig. 4E-H and Extended Data Fig. 4G).

Unlike WAPL depletion, singular RAD51 siRNA failed to rescue cohesion fatigue in 14 Gy irradiated cultures (Fig. 4F), and WAPL and RAD51 co-depletion imparted additive mitotic survival on G2 irradiated cells (Fig. 4G). WAPL and RAD51 depletion also differently affected the first mitotic duration under conditions of mitotic catastrophe (Fig. 4H). WAPL knockdown conferred a bi-modal mitosis length distribution where survival correlated with normal mitotic duration and death with mitotic arrest. In contrast, RAD51 depletion facilitated mitotic survival despite an extended mitotic duration, while also reducing the time of mitotic arrest to death (Fig. 4H). Further, ATR inhibition, which suppresses HR, also failed to rescue cohesion fatigue during mitotic catastrophe (Extended Data Fig. 4H).

### Suppressing HR rescues mitotic death at the cost of chromosomal instability

We live-imaged the chromosomal consequences of mitotic death escape in irradiated HeLa cultures expressing H2B-eGFP or mCherry-H2B (Fig. 5A and Supplementary Video 5). Chromosome misalignment was categorized as mild when one to a few chromosomes were present outside the metaphase plate, moderate when many chromosomes were dispersed but the metaphase plate remained visible, and severe when chromosomes either fail to form a metaphase plate or the metaphase plate becomes unstructured during mitotic arrest (Fig. 5A). Mitotic death was predominantly associated with severe misalignments, while cells that survived mitosis displayed mild to moderate defects (Fig. 5B). This was most evident in the first mitoses after IR (Fig. 5B). Even when irradiated cells displayed mild chromosome alignment defects, all surviving daughter cells presented chromosome segregation errors.

**Figure 5:**
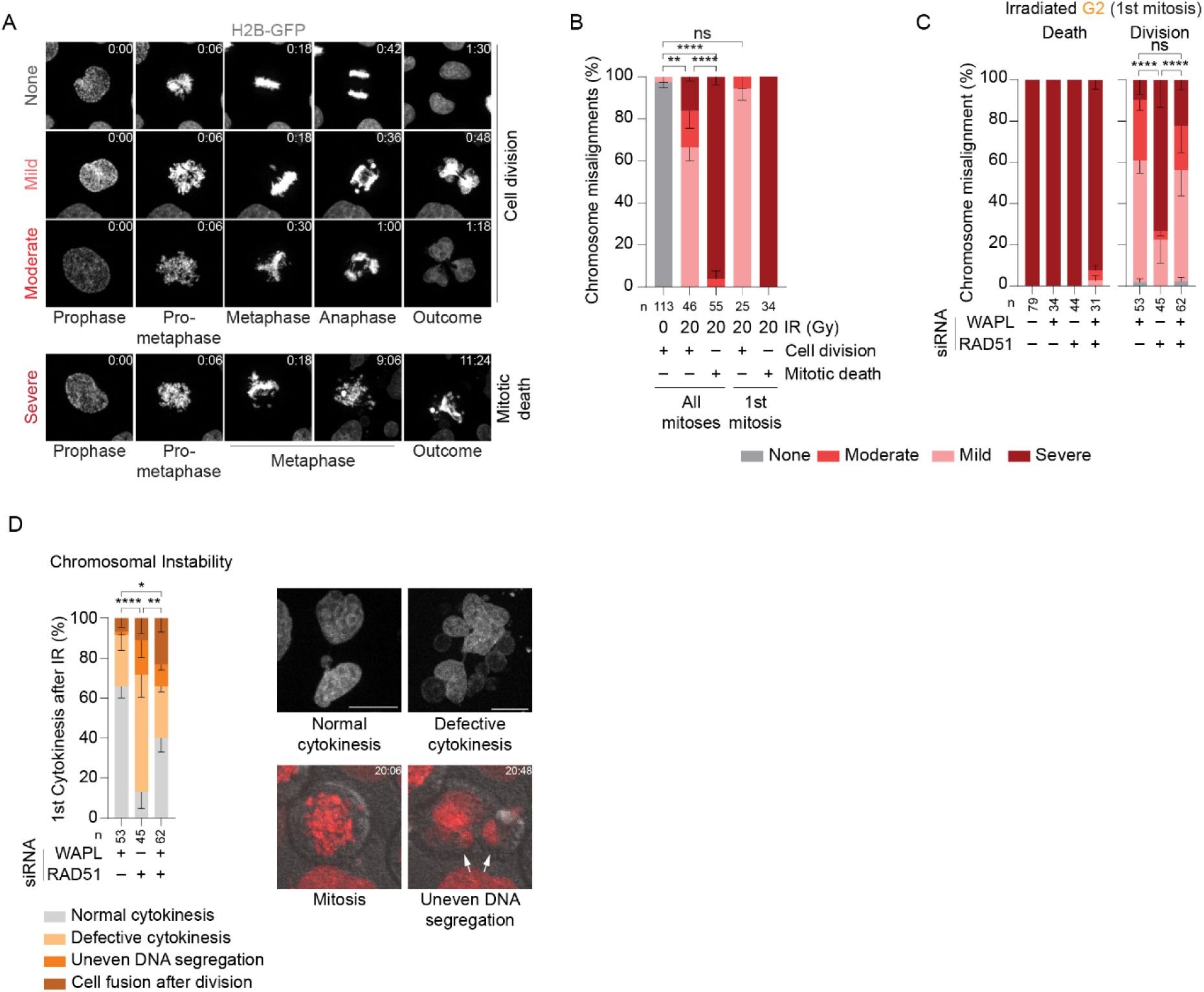
Targeting RAD51 promotes genomic instability in cells that escape mitotic death. **A)** Representative images from spinning-disk confocal live imaging of HeLa cells expressing H2B-eGFP following mock or 20 Gy IR. Time is hrs:min relative to prophase. **B)** Quantitation of the chromosome alignment shown in (A) for HeLa H2B-eGFP cells treated with mock or 20 Gy IR (mean ± SEM of 2 biological replicates; Fisher’s exact test). **C)** Chromosome alignment in the first mitosis of G2-enriched HeLa mCherry-H2B cultures transfected with the indicated siRNAs and treated with 14 Gy IR. Data are presented as a function of mitotic death or division (mean ± SEM of 3 biological replicates; Fisher’s exact test). **D)** Quantitation of cytokinesis defects in cells from (C) that successfully completed the first mitosis (mean ± SEM of 3 biological replicates; Fisher’s exact test). Representative images of normal and defective cytokinesis (right above), and uneven DNA segregation (right below) are shown. Time is hrs:min relative to start of the imaging following IR treatment. Where appropriate, ns = not significant, *p < 0.05, **p < 0.01, ****p < 0.0001. For panels B – D, n = number of cells quantified across all biological replicates.

Analysis of WAPL and/or RAD51 depleted mCherry-H2B HeLa cultures enriched for G2-phase (Extended Data Fig. 5A) and irradiated with 14 Gy IR revealed that nearly all cells undergoing immediate mitotic death exhibited severe chromosome alignment defects (Fig. 5C and Supplementary Video 6). Relative to singular RAD51 depletion, WAPL siRNA alone or in combination with RAD51 knockdown improved chromosome alignment in cells that survived the first mitosis after genome damage (Fig. 5C and Extended Data Fig. 5B). Within the population of cells that survived mitosis, the severity of mitotic chromosome mis-alignment directly correlated with subsequent chromosomal instability. Relative to singular RAD51 knockdown, WAPL depletion alone or with RAD51 siRNA reduced cytokinesis defects (Fig. 5D and Extended Data Fig. 5C). We interpret these data to signify that WAPL depletion rescues mitotic death by countervailing chromosomal misalignment and cohesion fatigue, whereas inhibiting HR enables mitotic exit despite these defects.

### Immediate mitotic death is executed through intrinsic apoptosis

To understand how DSB repair-mediated lethality is executed we used IncuCyte live imaging to quantify apoptosis through a cleaved Caspase-3/7 marker^49^ and cell proliferation through culture confluence (Fig. 6A and Extended Data Fig. 6A). Apoptotic death initiated within 8, 14, and 20 Gy irradiated cultures 32 to 36 hours post-treatment (Fig. 6A). A distinct second inflection in the apoptotic signal emerged 56 hours post-treatment in the 4 – 20 Gy treated cells. Proliferation was greatly reduced with 8 – 20 Gy dosages coinciding with early apoptotic death (Fig. 6A and Extended Data Fig. 6A).

**Figure 6:**
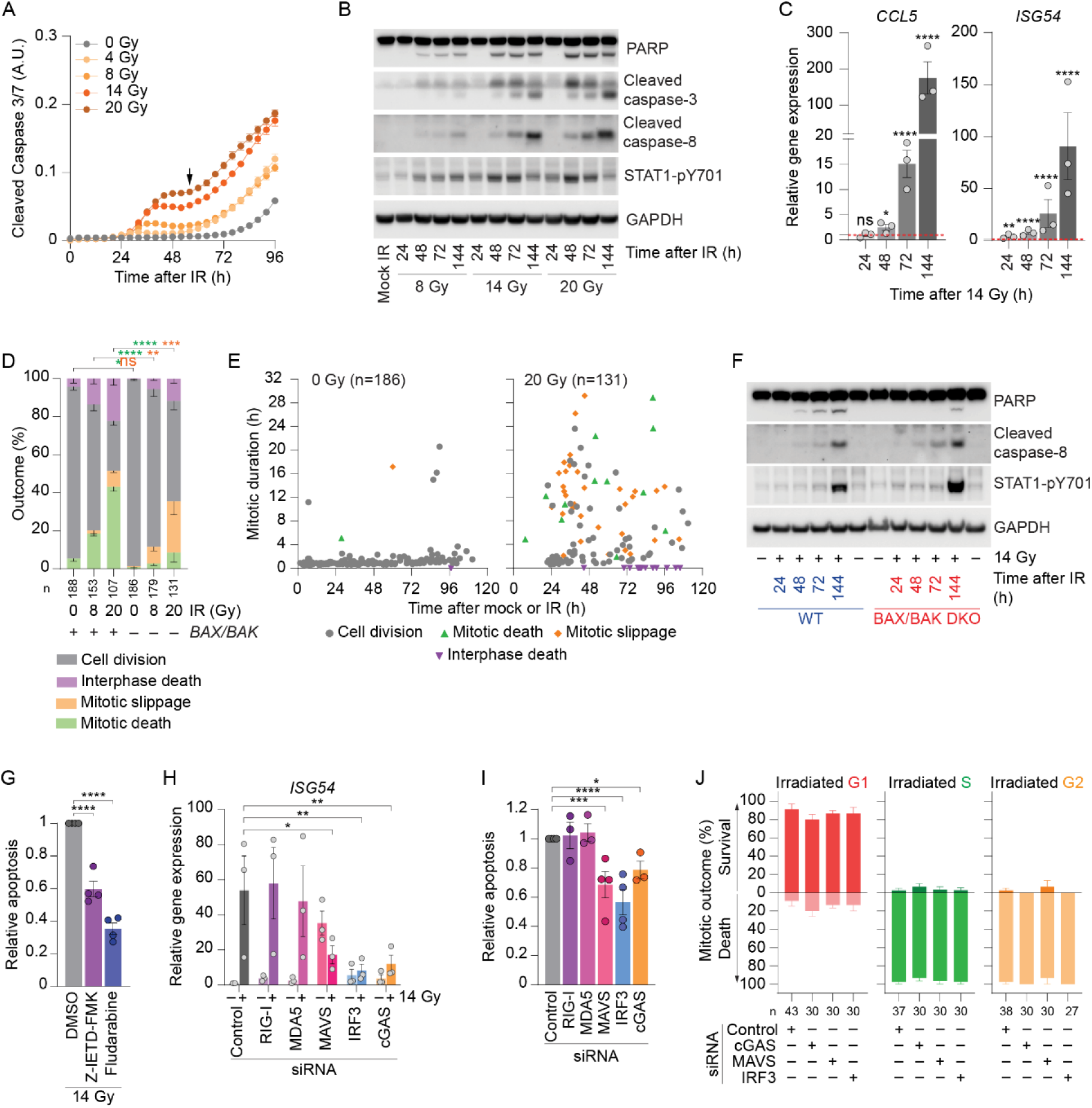
Distinct waves of intrinsic and extrinsic apoptosis during mitotic catastrophe. **A)** Apoptotic death in HeLa cultures measured through IncuCyte live imaging of fluorescent cleaved Caspase 3/7 reporter. A representative of three biological replicates is shown. (mean ± SEM, A.U. = arbitrary units). The arrow indicates second inflection of apoptotic signal. **B)** Western blots of whole cell extracts derived from HeLa cultures at the indicated time following mock or IR treatment. **C)** RT-qPCR measurement of interferon stimulated gene (ISG) expression in HeLa cells at the indicated times following 14 Gy IR (mean ± SEM of 3 biological replicates; RM one-way ANOVA with Fisher’s LSD test). Fold change in gene expression is relative to cells collected immediately prior to IR (T0), which is normalized to 1 and depicted as a red dotted line. **D)** Outcome measured by live imaging of HeLa parental, and *BAX* and *BAK* DKO cells following mock or IR treatment (mean ± SEM of ≥ 2 biological replicates; Fisher’s exact test). **E)** Multi-dimensional representation of the data from (D) graphed as in Fig. 1F. **F)** Western blots of whole cell extracts derived from HeLa parental, and *BAX* and *BAK* DKO cultures at the indicated time following mock or 14 Gy IR. **G)** Apoptosis measured as in (A) and shown in Fig. S6C in HeLa cultures treated with 14 Gy IR ± DMSO, 40µM Z-IETD-FMK, or 10µM Fludarabine (mean ± SEM; n = 4 biological replicates; RM one-way ANOVA with Fisher’s LSD test). **H)** *ISG54* expression measured by RT-qPCR in HeLa cultures treated with the indicated siRNAs ± 14 Gy IR (mean ± SEM; n = 3 biological replicates, RM one-way ANOVA with Fisher’s LSD test; depicted p values are relative to ctrl siRNA 14 Gy IR). Fold change is relative to non-irradiated cells treated with control siRNA and normalized to 1. **I)** Apoptosis in HeLa cultures treated with the indicated siRNAs and following 14 Gy IR, measured as in (G) (mean ± SEM; n ≥ 3 biological replicates; Mixed-effect one-way ANOVA with Fisher’s LSD test). **J)** First mitosis outcome following 14 Gy IR in 3F HeLa cells treated with the indicted siRNAs as a function of cell cycle phase at irradiation (mean ± SEM of 3 biological replicates). Where appropriate, ns = not significant, *p < 0.05, **p < 0.01, *** p <0.001, ****p < 0.0001. For panels D, E, and J, n = number of cells quantified across all biological replicates.

Intrinsic and extrinsic apoptosis share the effector Caspase-3/7, while the initiator Caspase-8 functions specifically in extrinsic apoptosis^1, 50, 51^. IR dose- and time-dependent Caspase-3 activation and PARP cleavage, known apoptotic hallmarks^52^, occurred contemporaneous with immediate mitotic and delayed lethality (Fig 6B, compare to Fig. 1F). Caspase-8 cleavage, however, was delayed relative to PARP and Caspase-3 (Fig. 6B) and peaked with delayed lethality 72 hours or more post-IR. The transcription factor STAT1 regulates Caspase-8 ^53, 54^ and pro-inflammatory ISG transcription^55, 56^. In agreement, activatory STAT1-Y701 phosphorylation preceded Caspase-8 activation, ISG transcription, and delayed lethality (Fig. 6A-C and Extended Data Fig. 6B).

Suppressing intrinsic apoptosis with *BAX* and *BAK* double knock out (*BAX BAK* DKO)^47, 57^ significantly reduced mitotic death, while interphase death remained prominent in cultures treated with 8 or 20 Gy IR (Fig. 6D, E) In agreement, PARP cleavage was deferred in irradiated *BAX BAK* DKO HeLa cells relative to parental cultures and instead overlapped with later Caspase-8 activation (Fig. 6F). STAT1-Y701 phosphorylation and Caspase-8 activation were also more robust in *BAX BAK* DKO cells relative to parental controls (Fig. 6F). HR-dependent mitotic death is therefore executed primarily through intrinsic apoptosis while NHEJ, MMEJ, and SSA-mediated delayed lethality can proceed independent of *BAX* and *BAK*.

### The innate immune response promotes delayed lethality

We probed for extrinsic death by treating irradiated HeLa cultures with STAT1 (10µM Fludarabine ^58^) or Caspase-8 (40µM Z-IETD-FMK ^59^) inhibitors, and measuring Caspase-3/7 cleavage. Both Fludarabine and Z-IETD-FMK significantly reduced apoptosis in 14 Gy irradiated cells at late time points (Fig. 6G and Extended Data Fig. 6C). Z-IETD-FMK and Fludarabine also reduced apoptosis in overly confluent mock irradiated cultures from 72 hours (Fig. 6G and Extended Data Fig. 6C). 14 Gy irradiated cultures, however, never approached confluence (Extended Data Fig. 6A), indicating the effects of STAT1 or Caspase-8 inhibition were independent of constrained growth.

Recognition of inappropriate cell body DNA by cGAS, or RNA by RIG-I or MDA5 that signal through the common mediator MAVS, promote STAT1 activation through nuclear translocation of IRF3^6, 60^ ^3^. Depleting IRF3, cGAS, or MAVS, but neither RIG-I nor MDA5 alone, suppressed ISG transcription and apoptosis in 14 Gy irradiated HeLa cultures (Fig. 6H, I and Extended Data Fig. 6D-G). Notably, depleting IRF3, cGAS, or MAVS failed to impact first cell cycle outcomes in 14 Gy irradiated 3F HeLa cells (Fig. 6J) indicating the innate immune response contributes to delayed lethality, but not immediate mitotic death.

### Targeting DSB repair modifies immunogenicity during mitotic catastrophe

Intrinsic apoptosis is non-immunogenic^61, 62^, suggesting that HR-dependent mitotic lethality is an immunologically cold response. Conversely, G1 irradiated cells survive the first mitosis with NHEJ-, MMEJ-, and/or SSA-mediated chromosome structural aberrations, and perish thereafter following innate immune response activation. This suggested a direct causal relationship between DNA repair pathway choice and immunogenicity.

Treating 14 Gy irradiated HeLa cultures with VE-822 suppressed HR and mitotic death (Fig. 3F and S3B), conferred enhanced delayed lethality (Fig. 3H), and significantly elevated ISG expression (Fig. 7A and Extended Data Fig. 7A). Suppressing mitotic death in 14 Gy IR treated cells with WAPL knockdown (Fig. 4G) also bestowed higher ISG expression (Fig. 7A and Extended Data Fig. 7A, B). Conversely, LIG4 or RAD52 depletion promoted mitotic death in 14 Gy irradiated cultures (Fig. 2E and Extended Data Fig. 2E) and suppressed ISG expression (Fig. 7A and Extended Data Fig. 7A, B). When HR was inhibited in irradiated RAD52 depleted cells with VE-822, this suppressed mitotic death (Fig. 3F) and restored the enhanced ISG expression (Fig. 7A and Extended Data Fig. 7A). Promoting HR therefore enhances non-immunogenic mitotic death and suppresses immunogenicity during mitotic catastrophe.

**Figure 7:**
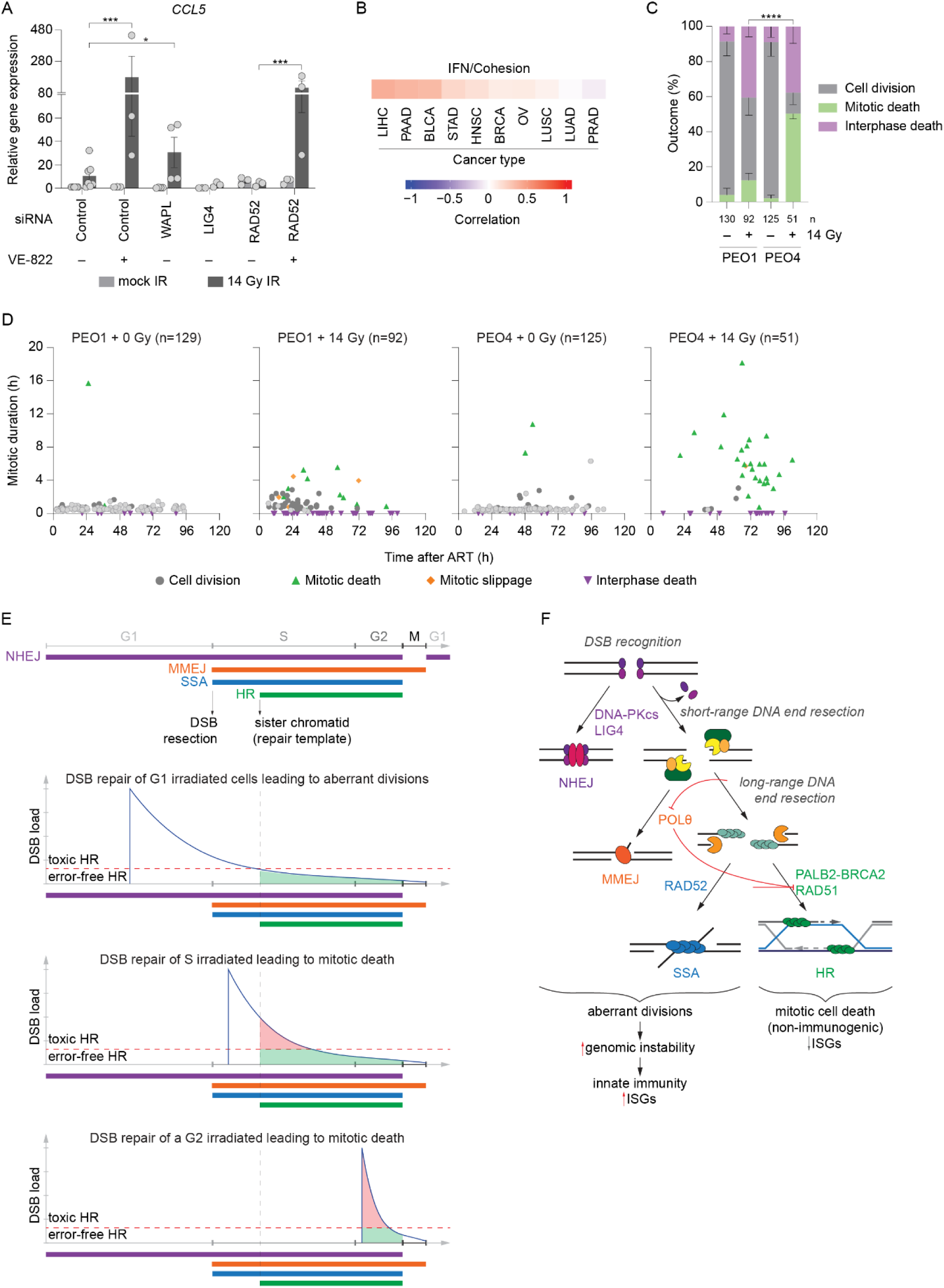
Mitotic catastrophe outcomes are pharmacologically tunable and influenced by disease relevant genetics. **A)** RT-qPCR measurement of *CCL5* expression in HeLa cultures treated with the indicated siRNAs ± 14 Gy IR and were indicated 0.2µM VE-822 (mean ± SEM; n ≥ 3 biological replicates; Mixed-effect one-way ANOVA with Fisher’s LSD test). Fold change in gene expression is relative to non-irradiated cells treated with control siRNA and normalized to 1. VE-822 was added one hour prior to mock or 14 Gy IR treatment. Asterisks denote comparisons between 14 Gy IR samples treated with the indicated siRNAs. **B)** Spearman correlation between IFN and cohesion signatures (as derived from mSigDB gene lists) in patient samples from The Cancer Genome Atlas. LIHC = liver hepatocellular carcinoma, PAAD = pancreatic adenocarcinoma, BLCA = bladder cancer, STAD = stomach adenocarcinoma, HNSC = head and neck squamous cell carcinoma, BRCA = breast cancer, OV = ovarian cancer, LUSC = lung squamous cell carcinoma, LUAD = lung adenocarcinoma, PRAD = prostate adenocarcinoma. **C)** Outcome measured through live imaging of PEO1 and PEO4 cultures ± 14 Gy IR (mean ± SEM from 2 biological replicates, n = number of cells images across both replicates; Fisher’s exact test). **D)** Multi-dimensional representation of the data from (C) depicted as described in Fig. 1F. **E)** A model of DSB mediated lethality. Activity of DSB repair pathways across cell cycle (top). Interpretation of repair kinetics following DSB induction in G1, S, or G2 phase cells (below) as a function of DSB load and cell cycle progression. A red dotted line indicates a theoretical DSB threshold that separates functional (green shading) and toxic (red shading) HR. **F)** Proposed decision tree of DSB repair pathway choice with corresponding outcome following ablative radiation. Where appropriate, *p < 0.05, *** p <0.001, ****p < 0.0001.

We interrogated exome and RNA-sequencing data from The Cancer Genome Atlas (TCGA) and found across >1000 samples from 10 cancer types a generally positive correlation between expression of genes in the cohesion pathway and interferon gene signatures (Fig. 7B). This suggests that enhanced cohesion in cancer cells with underlying genome instability facilitates more effective proliferation and immunogenicity, reminiscent of our observations of the impact of WAPL depletion on irradiated cells.

### *BRCA2* mutations suppress mitotic death

Mutations in HR pathways drive the etiology of diverse cancer types and are frequent in many tumors^63, 64^. Our data suggest tumors harboring such mutations are potentially resistant to immediate mitotic death during mitotic catastrophe. PEO1 and PEO4 are ovarian cancer lines derived from the same patient that respectively carry inactivating and rescuing *BRCA2* mutations^65^. Consistent with immediate mitotic lethality being HR-dependent, HR-incompetent PEO1 cells were resistant to mitotic arrest and death following 14 Gy IR, while the same IR dose effectively induced mitotic arrest and immediate mitotic death in the HR-competent PEO4 cells (Fig. 7C, D).

## DISCUSSION

In this study, we visualized the outcomes of individual cells following lethal genomic damage. Our data establish that the repair of DSBs controls the distinct modes of immediate mitotic death and delayed lethality encompassed within the spectrum of mitotic catastrophe. Moreover, the pivotal decision between mitotic death versus mitotic survival in the first cell cycle dictates if the innate immune response is suppressed or activated in response to genomic insult. These findings elucidate the previously ambiguous concept of mitotic catastrophe and provide a roadmap for enhancing immunogenicity following clinical induction of DSBs (Fig. 7E, F).

Lethal genomic insult led to pronounced outcome bifurcation dependent upon cell cycle regulated DSB repair. Immediate mitotic death required SAC-dependent mitotic arrest promoted by the upstream HR factors ATR, BRCA2, PALB2, and RAD51. Mitotic death was counteracted by the downstream HR components RTEL1 and SLX4, which respectively prevent and resolve dHJs contingent upon upstream HR activity. Depleting BRCA2 and/or PALB2 suppressed HR initiation and rescued immediate mitotic lethality in RTEL1 or SLX4 depleted cells. These results directly implicate dHJs as the central mediator of toxic HR. Conversely, the combined activity of NHEJ, MMEJ, and SSA prevented immediate mitotic death in G1-irradiated cells. Although NHEJ is conceptualized as the sole DSB repair pathway active in G1, emerging evidence suggests the potential for limited G1-phase HR^13^. Under conditions of RAD52 depletion, discernable RAD51 foci manifested in irradiated HeLa cells exhibiting G1 DNA content. While acknowledging the possibility of early S phase cell representation, the occurrence of RAD51 foci in G1/early-S coincided with conditions where HR underpinned the immediate mitotic lethality of G1 irradiated cells.

We propose that a temporal repair hierarchy, coupled with the cumulative load of DSBs, serves as a reliable predictor of mitotic catastrophe outcomes (Fig. 7E, F). During G1, NHEJ swiftly ligates severed DNA at the potential expense of multicentric or ring chromosomes. However, in the absence of p53 signaling, chromatin carrying damage can pass from G1 to early-S^66^. Should NHEJ fall short of rectifying all G1-induced DSBs, MMEJ and SSA could undertake S-phase repair^11, 67^ at the potential cost of radial chromosomes or sister chromatid fusions. HR becomes active later in S^8, 10^. If the combined activities of NHEJ, MMEJ, and SSA prove insufficient to fully rectify breaks induced during G1, HR may engage with the remaining lesions. When the count or nature of breaks entrusted to HR surpasses a critical threshold, the consequence is immediate mitotic death (Fig 7E). While the preponderance of G1 irradiated cells survived the first mitosis, we did observe a significant IR dose-dependent increase in the minority of G1 irradiated cells that experienced immediate mitotic death (Fig. 1G). This aligns with a higher likelihood of G1 damage evading the combined efforts of NHEJ, MMEJ, and SSA as the DSB burden escalates commensurate with the IR dose. In contrast, induction of DSBs during S or G2 readily triggers HR engagement. The result is conspicuous mitotic death in S/G2 cells irradiated with 2 Gy and nearly universal mitotic lethality following S/G2 irradiation with 8 Gy. (Fig. 7E). Promoting dHJ formation or retention through RTEL1 or SLX4 depletion further accentuated mitotic death in response to low dose IR.

Long-range DNA end resection inhibits MMEJ, directing DSB repair towards HR or SSA (Fig. 7F)^11, 68^. Previous findings highlight the competition between SSA and HR as DSB load escalates^41, 69^. In such scenarios, excessive resection fosters a preference for SSA over HR, and prompts the substitution of RAD51 with RAD52 at DSBs^41, 69^. This phenomenon likely accounts for our observation of RAD51 foci in irradiated G1/early S cells subsequent to RAD52 depletion. We anticipate that with RAD52 siRNA, breaks that should have been demarcated for SSA are instead loaded with RAD51 and trigger HR, ultimately leading to immediate mitotic death. Concurrent depletion of RAD51 and RAD52, however, suppresses HR and rescues mitotic death in the SSA compromised cells.

When irradiated cells complete the first cell cycle during mitotic catastrophe, chromosome aberrations resulting from NHEJ, MMEJ, and SSA activate the innate immune response and promote delayed lethality (Fig. 7F). This is consistent with mitotic progression being necessary for cytosolic nucleic acid generation following genome damage^3–6^. Previous findings implicate cGAS and its effector IRF3 in mitotic death^70^. However, we observed no effect on immediate mitotic death with siRNAs against cGAS, IRF3, or the common RNA sensing node MAVS. This is consistent with immediate mitotic death occurring independent of the innate immune response. Based on our collective findings, we propose to clarify the IR-induced phenotypes previously encompassed within mitotic catastrophe as HR-dependent immediate mitotic death, and first cell cycle survival and delayed innate immune response activation.

Genetic or pharmacological interventions targeting DSB repair exerted distinctive effects on the innate immune response during mitotic catastrophe. Depleting LIG4 or RAD52 magnified mitotic death and suppressed ISG transcription. In contrast, inhibiting ATR or depleting WAPL facilitated first cell cycle survival and amplified ISG expression. Notably, rescuing mitotic death with ATR inhibition produced a stronger innate immune response than WAPL depletion. We speculate this is because WAPL knockdown enhanced cohesion and conferred more appropriately structured daughter nuclei, consistent with the generation of fewer cytosolic nucleic acids. Supporting this notion, patient data revealed that enhanced cohesion corresponds with suppressed interferon signaling, suggestive of clinical implications that warrant further exploration. We interpret our findings to indicate that targeting NHEJ or SSA accentuates HR-dependent mitotic death through non-immunogenic intrinsic apoptosis. Removal of damaged cells during the first mitosis prevents generation of the cytosolic nucleic acids required for innate immune response activation. Consequently, suppressing HR fosters heightened immunogenicity by enhancing first cell cycle survival with chromosome segregation defects that active cGAS, MAVS, and IRF3.

The precise mechanisms by which toxic dHJs evade nuclease or dissolution activity, and maintain persistent SAC activation, remain unknown. A hypothesis we favor involves the formation of irregular dHJs between non-sister chromatids. The likelihood of such anomalous repair events should increase proportionally with the dose of DSBs. It is also conceivable that dHJ resolution activities are less efficient when dealing with intermediates formed between non-sister chromatids. Additionally, we anticipate WAPL depletion reinforces cohesion during S and G2 phases^71^, promoting HR between physically linked sister chromatids.

Within the spectrum of cancer interventions, radiation therapy (RT) stands as a prime candidate to leverage mitotic catastrophe for patient benefit. Recent advances in medical imaging have led to development of highly precise ablative radiotherapy (ART), clinically termed stereotactic ablative radiotherapy (SABR), stereotactic body radiation therapy (SBRT), or stereotactic radiosurgery (SRS))^72–74^. ART is a standard of care for localized and oligometastatic disease^75, 76^ that is administered in one to five high-dose fractions ranging from 8 to 21 Gy^77^. ART provides enhanced tumor control^78, 79^, and critically, can synergize with immunotherapy^80, 81^. Ongoing translational efforts aim to enhance the ART therapeutic efficacy by bolstering the immunological response ^82, 83^.

Our findings yield two prospective strategies to enhance ART therapeutic ratio. First, driving mitotic death in early-stage cancers carrying NHEJ, MMEJ or SSA mutations, or potentiated by combining ART with pharmacological inhibition of these repair pathways, holds potential to widen the therapeutic index between normal and malignant tissues. This conjecture may explain the observed synergistic effects of combining Polθ inhibitors and conventional fractionated RT^84^. For patients with metastatic disease, we anticipate that suppression of HR during ART could promote the innate immune response and enhance the efficacy of combinatorial immunotherapy. Additionally, our findings suggest that patients with HR-deficient cancers are more likely to manifest an immunological response to ART. A personalized therapeutic approach that predicts conventional or ablative RT response as a function of tumor genetics could yield therapeutic benefit. Current overall response rates to immunotherapy hover near 15%^85^. Promoting immunogenicity with ART under conditions of pharmacological or genetic HR suppression has the potential to benefit patients worldwide.

## METHODS

### Cell Culture

HeLa (human, female) cells were obtained from Megan Chircop (CMRI, Sydney, Australia). H2B-eGFP HeLa were previously created and characterized ^86^. *BAX* and *BAK* double knockout (DKO) HeLa cultures generated from a parental CCL-2 strain were a gift from David Huang (WEHI, Melbourne, Australia), previously generated and characterized elsewhere ^47^. HT1080 6TG (human, male) cells were a gift from Eric Stanbridge (University of California, Irvine, US). IMR90 E6E7 (human, female) cells were a gift from Jan Karlseder (Salk Institute, La Jolla US) and 2-color FUCCI (2F) IMR90 cells were created previously ^47^. HCT116 p53 KO (human, male) were obtained from Bert Vogelstein (Johns Hopkins University, Baltimore, US). T98G (human, male) and A549 (human, male) were provided by Roger Reddel (CMRI, Sydney, Australia). PEO1 (human, female) and PEO4 (human, female) cells were a gift from Anna DeFazio (WIMR, Sydney, Australia). Phoenix-AMPHO (human, female) were bought from ATCC. The identity and purity of all cell lines was verified by Cell Bank Australia using short tandem repeat (STR) profiling. All cultures were routinely mycoplasma tested and found to be negative (MycoAlert, LT07-118, Lonza). All lines except PEO1, PEO4, and Phoenix-AMPHO were cultured at 37 °C, 10% CO2, and 3% O2 in DMEM (Life Technologies) supplemented with 1% non-essential amino acids (Life Technologies) and 1% Glutamax (Life Technologies). HeLa, HT1080 6TG, HCT116 p53 KO, and their derivatives were supplemented with 10% bovine growth serum (Hyclone), while T98G, A549, IMR90, and their derivatives were supplemented with 10% fetal bovine serum (Life Technologies). PEO1 and PEO4 cells were cultured at 37 °C, 10% CO2, and 3% O2 in RPMI (Life Technologies) supplemented with 20mM HEPES (Sigma-Aldrich, H0887), 1% Glutamax, and 10% fetal bovine serum. Phoenix-AMPHO cells were cultured at 37 °C, 10% CO2, and 3% O2 in DMEM (Life Technologies) supplemented with 1% non-essential amino acids (Life Technologies) and 10% fetal bovine serum (Life Technologies). Cells were treated with the following compounds: dimethyl sulfoxide (DMSO; Sigma-Aldrich, D8418), 0.5 µM Reversine (Selleckchem, S7588), 40 µM Z-IETD-FMK (Selleckchem, S7314), 10 µM Fludarabine (Selleckchem, S1491), 0.5 µM NU7441 (Selleckchem, S2638), 10 µM 5-Ethynyl-2′-deoxyuridine (EdU; Sigma-Aldrich, 900584), 10 µM RO3306 (Sigma-Aldrich, SML0569), 0.2 µM VE-822 (Selleckchem, S7102), 250 nM colcemid (Gibco ThermoFisher, 15212012), and 2 mM thymidine (Sigma-Aldrich, T1895). Cells were respectively irradiated as a monolayer or in suspension using an X-RAD 320 (1 Gy/min; Precision X-Ray) or GammaCell 3000 Elan (1.59 Gy/min ± 0.07; Best Theratronics) irradiator.

### IncuCyte proliferation and apoptosis quantitation

Cultures were imaged using an IncuCyte live imaging system (Sartorius) at 37 °C, 10% CO2 and 3% O2, with images captured at 10x magnification every 2 or 4 hours for the experiment duration. Proliferation was calculated based on a confluence mask generated with the IncuCyte Zoom software (Sartorius, version 2019B). We detected apoptosis using the NucView488 reagent (Biotium, 10402), which is a Caspase-3/7 enzyme substrate that emits green fluorescence upon apoptotic cleavage. 1:1000 NucView488 ± DMSO, 40 µM Z-IETD-FMK, or 10 µM Fludarabine were added into FluoroBrite (Gibco ThermoFisher, A1896701) media ≥ 30 minutes before mock or actual radiation. Caspase-3/7 activity was calculated based on generated cell mask and expressed as a total integrated intensity (for pharmacological treatments), or apoptotic cell count under conditions that affected cell size and/or morphology (genetic knockdowns). Apoptosis, as a temporal function of Caspase-3/7 activity, was calculated as an area under the curve relative to DMSO-treated cultures and normalized to 1 using GraphPad Prism v9.3.1.

### Viral transduction and cell line generation

Three color (3F) FUCCI cells were created using the tFucci(CA)2/pCSII-EF (RDB15446) ^29^ construct kindly provided by Dr Hiroyuki Miyoshi (Keio University) through RIKEN BRC. HeLa Cas9 cells were generated with the lentiCas9-Blast plasmid ^87^ (Addgene plasmid #52962; http://n2t.net/addgene:52962; RRID: Addgene_52962), a gift from Feng Zhang. High titer 3rd generation lentivectors harboring tFucci(CA)2/pCSII-EF or lentiCas9-Blast constructs were produced in the CMRI Vector and Genome Engineering Facility (Sydney, NSW, Australia) and cell cultures transduced for 24 (HeLa Cas9) or 48 (3F HeLa) hours with an MOI of 3 in media supplemented with 4 µg/ml of polybrene. HeLa Cas9 cultures were selected with 10 µg/ml Blasticidin (Sigma-Aldrich, SBR00022) for 6 days. 3F HeLa cells were cultured for 6 days and sorted at the Westmead Institute for Medical Research Flow Cytometry Facility (Sydney, NSW, Australia) for mCherry expression. 3F cultures were expanded for six days in 1µg/ml of Normocin (Invivogen, ant-nr-1) supplemented media and appropriate cell cycle-dependent colorimetric signaling verified by live imaging.

pLXSN H2B-mCherry plasmid was a gift from Laure Crabbe, The Centre for Integrative Biology, Toulouse, France ^88^. We moved mCherry from the C-to the N-terminus of histone variant H2B through PCR amplification of H2B, mCherry, and pLSXN backbone (Supplementary Table 1), followed by infusion cloning (Takara, 638911) according to the manufacture’s protocol. Retroviral vectors were created by transfecting Phoenix-AMPHO cells with pLXSN mCherry-H2B using Lipofectamine 3000 (Invitrogen Thermo Fisher Scientific, L3000015)) according to manufacturer’s instructions. Viral supernatant was used to infect HeLa cells and stable mCherry-H2B cultures were selected by treatment with 600 µg/ml G418 (Sigma-Aldrich, 4727878001). The fluorescent mCherry signal was confirmed by live imaging.

### Live cell imaging

Live cell imaging was performed in the CMRI Telomere Analysis Center (ATAC) as described previously ^47^ with the indicated modifications. All experiments were carried out at 37 °C, 10% CO2 and 3% O2 with imaging commencing as soon as possible after mock or actual irradiation. Events within the first 45 – 60 min following IR were not recorded due to the time required for sample transport from the irradiator and imaging setup. Images were captured every six minutes for the experiment duration. Differential interference contrast (DIC) with or without accompanying epifluorescence was acquired on a ZEISS Cell Observer inverted wide field microscope fitted with a 20 × 0.8 NA air objective and Axiocam 506 monochromatic camera (ZEISS) using ZEN software (ZEISS). A ZEISS HXP 120C mercury short-arc lamp and compatible filter cubes were used to generate fluorescent images. Chromosome dynamics in mCherry-H2B cells was captured using a ZEISS Cell Observer SD spinning disc confocal microscope fitted with 40 × 1.3 NA air objective, a 561 nm 50 mW solid state excitation laser (30% excitation power, 1 × 1 binning, EM gain of 300) with appropriate filters, and an Evolve Delta camera (Photometrics) using ZEN software (ZEISS). H2B-eGFP HeLa cultures were imaged on a Cell Voyager confocal microscope equipped with 40 × 0.95 NA objective (Olympus), 488nm laser (10 – 20% excitation power, 1 × 1 binning, 65% gain) with compatible filter cubes, and a high sensitivity EMCCD camera using CV1000 software (Yokogawa). 11 – 13 Z-stacks were taken at 1µm increments and outcomes were scored manually. For DIC microscopy, outcomes were assigned based on morphological features. Mitotic duration was calculated from the frame preceding nuclear envelope breakdown until cytokinesis and complete separation of genetic material, typically two or three frames after anaphase onset, or first signs of mitotic cell death (cell membrane blebbing). For FUCCI cultures, cell cycle duration was scored by eye based on fluorescence ^29, 34^. For mCherry-H2B and H2B-eGFP HeLa cultures, chromosome misalignments were scored by eye. All live cell imaging analyses were performed using ZEN or CV1000 software.

### Western blotting

Preparation of whole cell extracts and western blots were performed as described previously ^89^ and luminescence was visualized on an LAS 4000 imager (Fujifilm). Primary antibodies used in the study: PARP (46D11, Signaling Technology, 9532, 1:1000), Cleaved Caspase-3 (Asp175, Cell Signaling Technology, 9661, 1:500), Cleaved Caspase-8 (Asp384, 11G10, Cell Signaling Technology, 9748, 1:1000), STAT1-pY701 (58D6, Cell Signaling Technology, 9167, 1:1000), GAPDH (D16H11, Cell Signaling Technology, 5174, 1:5000), DNA-PKcs (Y393, Abcam, ab32566, 1:1000), LIG4 (EPR16531, Abcam, ab193353, 1:1000), β-Actin (AC-15, Sigma-Aldrich, A5441, 1:10,000), RAD51 (14B4. Novus, NB100-148, 1:500), RAD52 (F-7, Santa Cruz, sc-365341, 1:500), POLθ (Thermo Fisher Scientific, PA5-69577, 1:250), Histone H2B (V119, Cell Signaling Technology, 8135, 1:1000), BRCA2 (Ab-1, clone 2B, Sigma-Aldrich, OP95, 1:1000), PALB2 (Bethyl, A301-246A, 1:1000), vinculin (hVIN-1, Sigma-Aldrich, V9131, 1:10,000), WAPL (A-7, Santa Cruz, sc-365189, 1:1000), CHK1 (2G1D5, Cell Signaling Technology, 2360, 1:1000), CKH1-pS345 (133D3, Cell Signaling Technology, 2348, 1:1000), and β-Tubulin (Abcam, ab6046; 1:500). Secondary antibodies used in the study: Goat Anti-Mouse HRP (Dako, P0447, 1:5000 – 1:20,000) and Goat Anti-Rabbit HRP (Dako, P0448, 1:5000 – 1:20,000).

### RT-qPCR

Total RNA was isolated from cell pellets collected by trypsinization using the RNeasy Mini Kit (Qiagen, 74104) according to the manufacturer’s instructions. Genomic DNA was eliminated by on-column digestion for 30 minutes at 37 °C with DNase I (Thermo Fisher Scientific, EN0521). 500 ng of total RNA as determined by NanoDrop spectrophotometer (Thermo Fisher Scientific, ND-1000) was reverse-transcribed using the SensiFast cDNA Synthesis Kit (Meridian Bioscience, BIO-65054) according to the manufacturer’s instructions, and diluted 5 times. qPCR was performed using 2 µl of diluted cDNA, 200nM gene-specific primers designed to span exon-exon junctions whenever possible (Supplementary Table 2), and PowerUp SYBR Green Master Mix (Thermo Fisher Scientific, A25742). A total volume of 10µl per reaction was run on a QuantStudio 6 Flex Real-Time PCR System (Applied Biosystems Thermo Fisher Scientific, 4485691) for 45 cycles (2 minutes at 50 °C, 2 minutes at 95 °C, then cycling for 1 second at 95 °C and 30 seconds at 60 °C). Relative gene expression was normalized to GAPDH, and fold change (FC) was calculated as 2^-ΔΔCt^. Statistical analysis was performed using ΔΔCt values.

### siRNA transfection

Transient gene knockdown was achieved by using individual or pooled siRNA (Supplementary Table 3) at the final concentration of 10nM. Reverse siRNA transfection was performed 72 hours prior to irradiation using Lipofectamine RNAiMAX (Invitrogen Thermo Fisher Scientific, 13778150) according to the manufacturer’s instructions.

### CRISPR-mediated repair assay

We measured NHEJ and MMEJ repair of a genomic loci through Tracking Indels by Decomposition (TIDE) using a gRNA targeting a single genomic location (LBR2) known to be repaired by NHEJ and MMEJ with different kinetics ^36^. The LBR2 Alt-R gRNA was generated by Integrated DNA technologies (IDT) (GCCGATGGTGAAGTGGTAAG). TracrRNA:gRNA duplex was transfected according to manufacturer’s protocol. Briefly, tracrRNA and gRNA were hybridized at 95 °C for 5 minutes in a 1:1 ratio. After hybridization, the mixture was combined with RNAiMAX diluted in Opti-MEM (Gibco Thermo Fisher Scientific, 11058021), and incubated at room temperature for 20 minutes. 72 hours post-transfection, cells were lysed with Direct-PCR lysis (Viagen, Cat. 301-C) and incubated overnight with 0.2 mg/ml of Proteinase K (Roche, 03115801001) at 55 °C. A PCR spanning the LBR2 CRISPR site was performed using MyTaq Red Mix (Meridian, BIO-25043), and the PCR products were subjected to Sanger Sequencing. Editing efficiency and indel pattern of the LBR2 gRNA upon DNA-PKcs inhibition and POLθ depletion through TIDE ^90^. MMEJ and NHEJ ratios were calculated using the following formulas: frequency -7 deletion/((frequency +1 insertion) + (frequency -7 deletion)) and frequency +1 insertion/((frequency +1 insertion) + (frequency -7 deletion)), respectively.

### DSB repair reporter cell lines

The DSB repair reporter vectors pDRGFP (Addgene plasmid #26475; http://n2t.net/addgene:26475; RRID: Addgene_26475) and hprtSAGFP (Addgene plasmid #41594; http://n2t.net/addgene:41594; RRID: Addgene_41594), were a gift from Maria Jasin. DSB reporter cell lines were generated by transfection of HeLa cells with reporter constructs using Lipofectamine 3000 (Invitrogen Thermo Fisher Scientific, L3000015) according to the manufacturer’s instructions. Stable integration of reporter constructs (containing a puromycin resistance gene) was confirmed by selection with puromycin (1µg/ml), which was added 48 hours after transfection and maintained in cell media for 2 weeks.

To measure DSB repair, reporter cell lines were transfected with pCBASce (Addgene plasmid #26477; http://n2t.net/addgene:26477; RRID: Addgene_26477, a gift from Maria Jasin) plasmid using Lipofectamine 3000 according to the manufacturer’s instructions, resulting in production of the endonuclease I-SceI. I-SceI induces a DSB within stably integrated reporter constructs, and subsequent GFP expression acts as a measure of DSB repair ^38, 39^. Reporter cells were transfected in parallel with pCAGGS-mCherry (a gift from Phil Sharp; Addgene plasmid #41583; http://n2t.net/addgene:41583; RRID: Addgene_41583) as an internal control to estimate transfection efficiency and calculate relative DSB repair activity. At 48 hours post transfection, cells were collected and GFP and mCherry expression quantified on FACSCantoII (BD Biosciences) using FACSDiva 6.1.3 software, and analyzed using FlowJo v10.8.0. Relative DSB repair capacity was calculated as a ratio of the GFP positive cells (%) to mCherry positive cells (%) and normalized to a control condition (equal 1).

### RAD51 foci labelling

72 hours after siRNA transfection, cells grown on sterile glass coverslips were treated with DMSO or 0.2 µM VE-822, and simultaneously pulse-labelled with 10 µM EdU for 1 hour. Cells were either fixed immediately (4% paraformaldehyde in PBS for 10 minutes at 4 °C), or irradiated and maintained for 2 hours with 10 µM RO-3306 and DMSO or 0.2 µM VE-822 before fixation. All wash steps were performed in PBS with 0.1% Tween. Samples were washed three times, permeabilized with 0.5% Triton X-100 (in PBST) for 5 minutes, washed three times again, and blocked with 2% BSA (in PBST) for 2 hours. Samples were incubated overnight at 4 °C with primary anti-RAD51 antibody (Calbiochem, PC130, 1:500 in 2% BSA), then washed five times. All subsequent steps were performed in the dark. Coverslips were incubated with a click-iT EdU solution (10 μM 6-carboxyfluorescein-TEG azide (Berry and Associates, FF 6110), 10 mM sodium L-ascorbate (Sigma Aldrich, A4034), 2 mM Copper (II) sulphate (Sigma Aldrich, 451657 in PBS) for 30 minutes. Cells were washed five times in 1% BSA (in PBST) before staining with Alexa Fluor 568-conjugated secondary antibody (Invitrogen, A11036, 1:500 in 2% BSA) for 1.5 hours. Cells were washed three times, incubated with DAPI (Sigma-Aldrich, 10236276001) for 5 minutes (1:5000 in PBST), washed 1x, rinsed twice with milliQ H_2_O, and sequentially dehydrated in 70%, 90%, and 100% ethanol for 3 minutes each. Air-dried coverslips were mounted with Prolong Gold Antifade Mountant (Invitrogen, Thermo Fisher Scientific, P36934), and cured in the dark for at least 24 hours. Fixed cell microscopy images were acquired with a ZEISS AxioImager Z.2 microscope fitted with a 40 × 1.3 NA oil objective, Axiocam 506 monochromatic camera using ZEN software. 19 Z-stacks for each channel were taken at 0.24 µm increments.

### Image analysis of RAD51 foci

The extended depth of focus function in ZEN blue software v2.3 was used to make maximum projections of each image, then .CZI files were converted to a single channel .TIFF files and imported into CellProfiler v4.2.1. Using custom image analysis pipelines, we pre-processed images using the CorrectIllumination and EnhanceOrSuppress feature functions. Nuclei and subnuclear foci were identified using the IdentifyPrimaryObjects functions with either Minimum Cross-Entropy or Otsu thresholding. Foci were associated with their respective nuclei using the Relate function, and foci intensities were measured using the MeasureObjectIntensity function. Data were plotted in GraphPad Prism v9.3.1 using a double gradient colourmap for the Multiple Variables plots.

### Cytogenetic chromosome spreads and fluorescent hybridization in situ (FISH)

Cytogenetic chromosome spreads were performed as previously described ^91^. Briefly, cells were arrested in mitosis with 250 nM colcemid (Gibco Thermo Fisher Scientific, 15212012) for 1 hour prior to fixation with methanol and acetic acid (3:1). Cells were subsequently dropped onto glass slides, fixed with 2% paraformaldehyde, and dehydrated in a graded ethanol series (70%, 90%, 100%) prior to performing fluorescent in situ hybridization (FISH). For FISH, cells were denatured at 80 °C for 10 minutes then hybridized in the dark overnight with peptide nucleic acid (PNA) probes against telomeric (Alexa488-OO-ccctaaccctaaccctaa; Panagene, F1004) and centromeric (Alexa647-OO-aaactagacagaagcatt-KK; Panagene, F3015) sequences. All subsequent steps were performed in the dark. Slides were washed with PNA wash A (70% formamide and 10mM Tris (pH 7.5)), followed by PNA wash B (50mM Tris (pH 7.5), 150mM NaCl and 0.8% Tween20), with DAPI added in the latter wash as a counterstain. Slides were then mounted using Prolong Gold Antifade, and allowed to set overnight before imaging. Metaphase searching and image acquisition was performed as previously described ^66^ using automated MetaSystems imaging platform on a ZEISS AxioImager Z.2 microscope fitted with a 63 × 1.4 NA oil objective, appropriate filter cubes, and a CoolCube1 camera (MetaSystems).

### Image analysis of cytogenetic chromosome spreads

Metaphase images were analyzed using ISIS software (MetaSystems). For structural and cohesion analysis, images were scored by eye. Metaphases containing a single chromatid-type fusion or radial chromosome were scored as positive for those events. Cohesion status within a single metaphase was typically consistent across all chromosomes. However, when multiple cohesion fatigue phenotypes were present within one metaphase, cohesion status was assigned as previously described ^47^. Briefly, cohesion status was considered normal, if < 3 chromosomes exhibited aberrant cohesion, mild if ≥ 3 chromosomes exhibited centromeric cohesion loss but retained cohesion between chromosomal arms, moderate if ≥ 3 chromosomes exhibited centromeric cohesion loss together with separation of adjacent chromosomal regions, and severe, if ≥ 3 chromosomes displayed complete sister chromatid separation.

### Cell cycle synchronization

Cells were synchronized at G1/S by a double thymidine block. Briefly, cells were treated with 2 mM thymidine for 16 – 18 hours, released for 6 – 8 hours, and treated again with 2 mM thymidine for an additional 16 – 18 hours. Following a release from the second thymidine treatment, cells were collected at different times and cell cycle profiles were determined by flow cytometry or by FUCCI coloration.

### Cell cycle analysis by flow cytometry

Cells were fixed in ice-cold 70% ethanol and immediately processed or stored overnight at – 20 °C. Cells were pelleted, washed in PBS, and resuspended in a propidium iodide (1mg/ml) and RNase A (0.5mg/ml; Qiagen, 1007885) staining solution. Cell cycle distribution was quantified on FACSCantoII (BD Biosciences) using FACSDiva 6.1.3 software and analyzed using FlowJo v10.8.0.

### TCGA analysis

Gene expression and somatic mutation data were obtained for samples across cancer types from the Cancer Genome Atlas (TCGA) accessed through the Broad Institute portal Firebrowse (http://firebrowse.org/). RSEM normalized, log2(x+1) transformed gene expression values were used for gene signature evaluation (signatures were taken as lists of genes from the Broad Institute database MSigDB), and median of signature genes’ expression was taken as the signature score for each patient sample. Spearman correlations were computed using the ppcor package in R between pairs of signature scores across TP53 mutated samples (non-synonymous, non-intronic, and not in an intergenic region) for a given cancer type. Correlations were only calculated if both groups had at least 10 non-NA values, and only if at least 10 samples were present with paired mutation and gene expression information for the cancer type.

### Statistical analysis

Statistical analysis was performed using GraphPad Prism v9.3.1. Statistical methods, number of replicates (n), and error bars are described in figure legends. Throughout the paper, ∗ indicates p < 0.05, ∗∗ indicates p < 0.01, ∗∗∗ indicates p < 0.005, ∗∗∗∗ indicates p < 0.001. Violin plots represent a cumulative distribution of data points from all replicates per condition. Representative data, whenever shown, are characteristic of similar results from at least two independent biological replicates. Figures were prepared using Adobe Photoshop and Illustrator.

## DATA AVALIBILY

This paper analyzes existing, publicly available data that can be located here: http://firebrowse.org/ (LIHC, PAAD, BLCA, STAD, HNSC, BRCA, OV, LUSC, LUAD, PRAD). All original code has been deposited at Zenodo and is publicly available here: https://github.com/andrewdhawan/miRNA_hallmarks_of_cancer/blob/master/1_linear_model_miRNA_mRNA.R (DOI: 10.5281/zenodo.1453559) (DOI: 10.5281/zenodo.1453559). The remaining datasets generated and/or analyzed during the current study are available from H.E.G. and A.J.C. upon reasonable request. Blot and quantitative source data will be provided should the manuscript proceed to resubmission. The 3F and 2F cell lines used in this study are subject to a material transfer agreement between the Riken BioResource Research Center, A.J.C., and the Children’s Medical Research Institute.

## ACKNOWLEDGEMENTS

Will Hughes, Scott Page, and The CMRI ACRF Telomere Analysis Center supported by the Australian Cancer Research Foundation, are thanked for microscopy infrastructure and support. The Westmead Institute of Medical Research Flow Cytometry Centre, supported by the Cancer Council NSW and the Australian NHMRC, are thanked for cell sorting. Anna De Fazio is thanked for cell lines. Members of the Cesare, Gee, Hau and Pickett laboratories are thanked for their critical feedback. A.G.M. is supported by a Human Frontiers Science Program fellowship, and A.J.C. by an Australian Research Council Future Fellowship (FT210100858). This project was supported by grants from the Australian NHMRC grants to A.J.C. (1106241), and A.J.C. and H.G. (2004430); an Australian Medical Research Future Fund award to H.P. and A.J.C. (2007488); Westmead Charitable Trust Early Clinician-Researcher Grant to H.G.; and funding from the Sydney West Radiation Oncology Network to H.G. and A.J.C..

## AUTHOR CONTRIBUTIONS

Conceptualization, R.S., H.E.G. and A.J.C.; Methodology, R.S., S.C., L.F., A.G.M., A.D., C.N., and S.G.P; Software, A.D.; Investigation, R.S., S.C., L.F., A.G.M., L.C., and C.N., Formal analysis, R.S., S.C., L.F., A.G.M., and A.D., Resources, H.A.P, E.H., H.E.G. and A.J.C., Funding Acquisition, A.G.M, H.A.P, E.H., H.E.G. and A.J.C., Writing – original draft, R.S., H.E.G. and A.J.C, with editorial contribution from S.C., L.F., C.N., and H.A.P.; Visualization, R.S., S.C., L.F., A.G.M., A.D., and A.J.C, Project Administration, R.S., H.G., and A.J.C.; Supervision, E.H., H.A.P., H.E.G. and A.J.C.

## COMPETING INTERESTS

H.A.P. is a co-founder and shareholder of Tessellate Bio. The remaining authors declare no competing interests.

**Correspondence and requests for materials** should be addressed to H.E.G. and A.J.C.

---

**Extended Data Figure 1:**
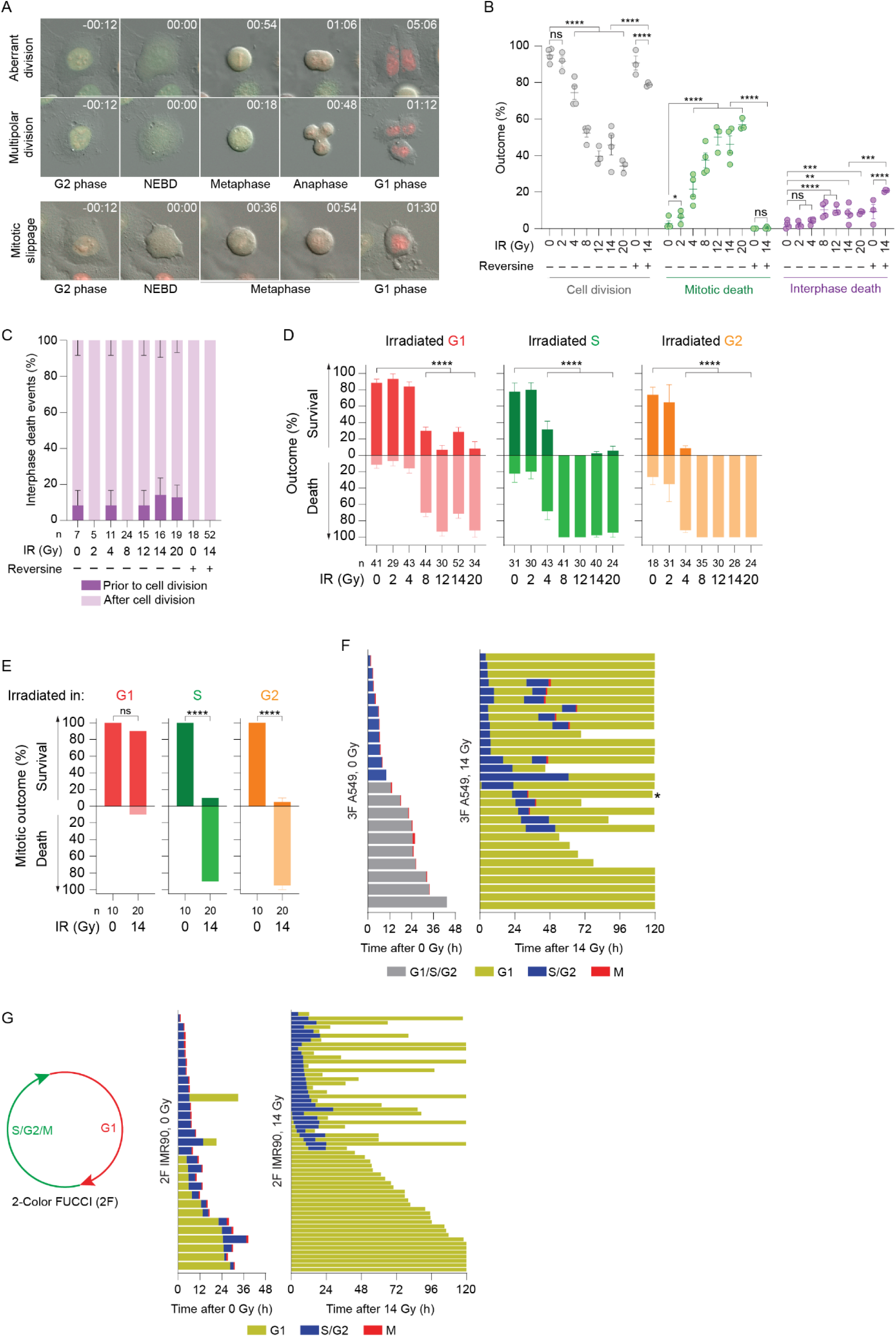
Double strand breaks confer distinct cell cycle-regulated outcomes. **A)** Representative images of aberrant mitotic outcomes from live imaging of 3F HeLa cultures. Time is hrs:min relative to NEBD. Scale bar = 50 µm. **B)** Means of the individual biological replicates from Fig. 1C (mean ± SEM; n ≥ 3 biological replicates; Fisher’s exact test). **C)** Quantitation of interphase lethality in Fig. 1C relative to first cell division following mock or IR treatment (mean ± SEM of ≥ 3 biological replicates). **D)** Outcomes across the entire 120-hour live imaging experiment as a function of cell cycle phase at irradiation (mean ± SEM of ≥ 3 biological replicates; Fisher’s exact test). This differs from Fig. 1G where only the first cell cycle was considered. **E)** First mitosis outcome measured by live imaging of 3F HT1080 6TG cultures as a function of cell cycle phase at irradiation (mean ± SEM of 2 biological replicates; Fisher’s exact test). **F, G)** Cell fate map of 3F A549 (G) and 2F IMR90 (H) cultures following mock or IR treatment. Each line represents a single cell as it progresses through cell cycle, relative to irradiation at T0. Track length corresponds to the time a cell spent in each cell cycle phase. Cell cycle phases are color coded as indicated in the legend. For mock-treated cultures, only the first cycle was recorded. For irradiated cultures, cells were tracked for as long as possible within 120 hours of imaging. The asterisk depicts a cell death event that occurred in the indicated cell cycle phase. All other tracks shorter than 120 hours represent cells that moved outside the field of view during the imaging. Data from one (3F A549) or two (2F IMR90) biological replicates are compiled into a single graph. The schematic of 2-color FUCCI is shown in H^34^. Where appropriate, ns = not significant, *p < 0.05, **p < 0.01, *** p <0.001, ****p < 0.0001. For panels C, E, F, and I, n = number of cells quantified across all biological replicates.

**Extended Data Figure 2:**
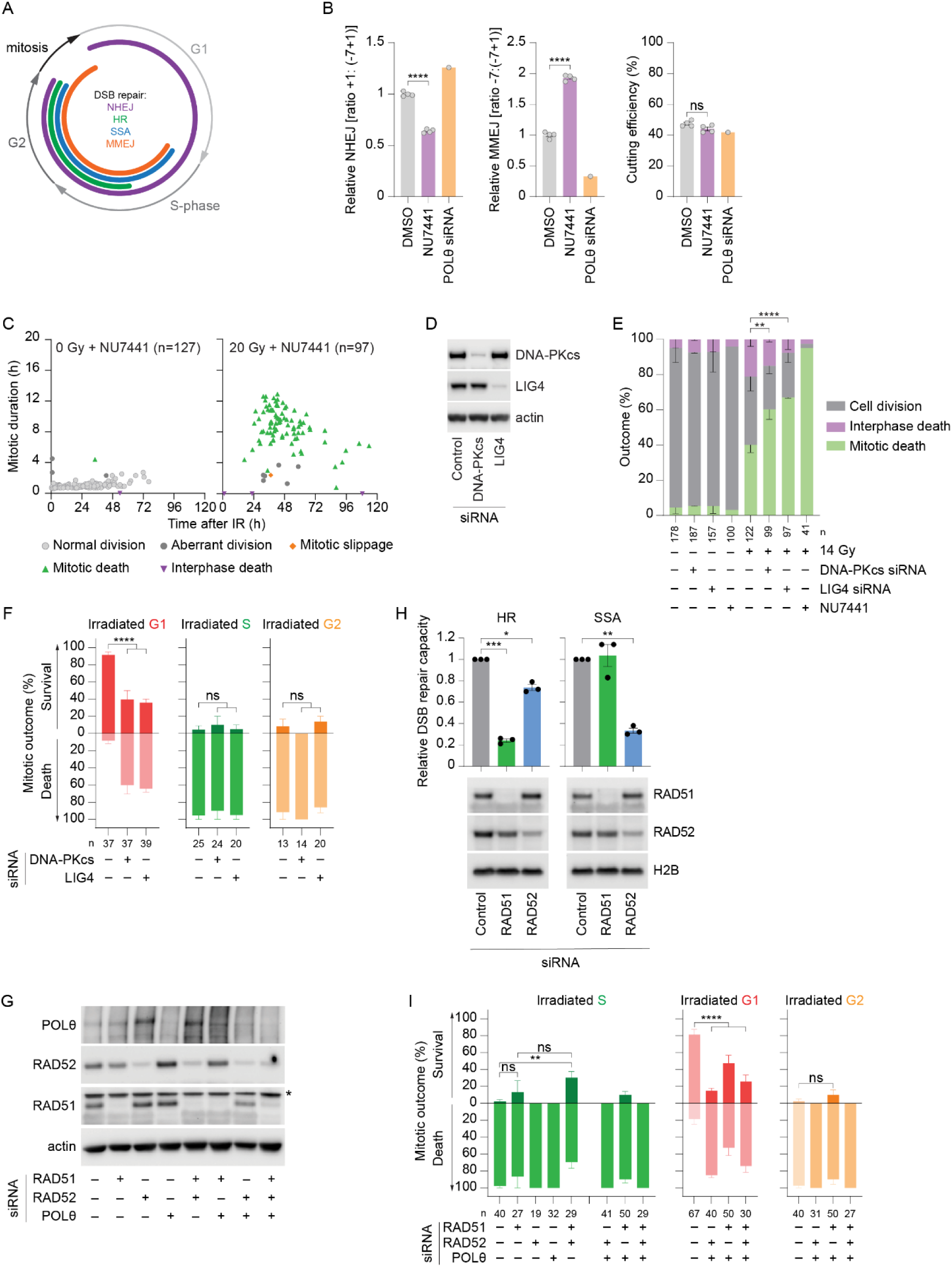
Immediate mitotic death is promoted by RAD51 and suppressed by DNA-PKcs, LIG4, RAD52, and Polθ. **A)** Diagram of active DSB repair pathways relative to cell cycle phase. **B)** TIDE measurement of relative NHEJ and MMEJ repair of a Cas9-induced DSB in HeLa cells. NHEJ and MMEJ activity were calculated on the respective frequency of +1 and -7 indels and normalized to 1 for DMSO-treated samples. Cas9 cutting efficiency is shown on right (mean ± SEM; n = 4 biological replicates for DMSO and NU7441, and one for POLθ siRNA; unpaired t-test). **C)** Multi-dimensional representation of data from Fig. 2A shown as described for Fig. 1F. **D)** Western blots of whole cell extracts derived from 3F HeLa cultures transfected with the indicated siRNAs. **E)** Outcomes measured by live imaging of 3F HeLa cultures transfected with the indicated siRNAs or 0.5µM NU7441 ± 14 Gy IR treatment (mean ± SEM of 2 biological replicates except for NU7441 treated cells (1 biological replicate) which served as a positive control; Fisher’s exact test; p values for mitotic death are depicted). **F)** Data from (E) classified by first mitosis outcome as a function of cell cycle phase at irradiation (mean ± SEM or 2 biological replicates, Fisher’s exact test). **G)** Western blot of whole cell extracts derived from 3F HeLa cultures transfected with the indicated siRNAs (*denotes a non-specific band). **H)** HR- or SSA-mediated repair of DSB reporters in siRNA and I-SceI transfected HeLa cells as measured by flow cytometry (mean ± SEM; n = 3 biological replicates; paired t-test). Western blots of whole cell extracts derived from the cultures used in repair quantitation are shown below. **I)** Fist mitosis outcome in 3F HeLa cultures transfected with the indicated siRNAs and treated with 14 Gy IR (mean ± SEM of ≥ 3 biological replicates). The data 14 Gy samples without siRNA for the G1 and G2 phase irradiated cells are the same as shown in Fig. 2E. Where appropriate, ns = not significant, *p < 0.05, **p < 0.01, *** p <0.001, ****p < 0.0001. For panels C, E, F, and I, n = number of cells quantified across all biological replicates.

**Extended Data Figure 3:**
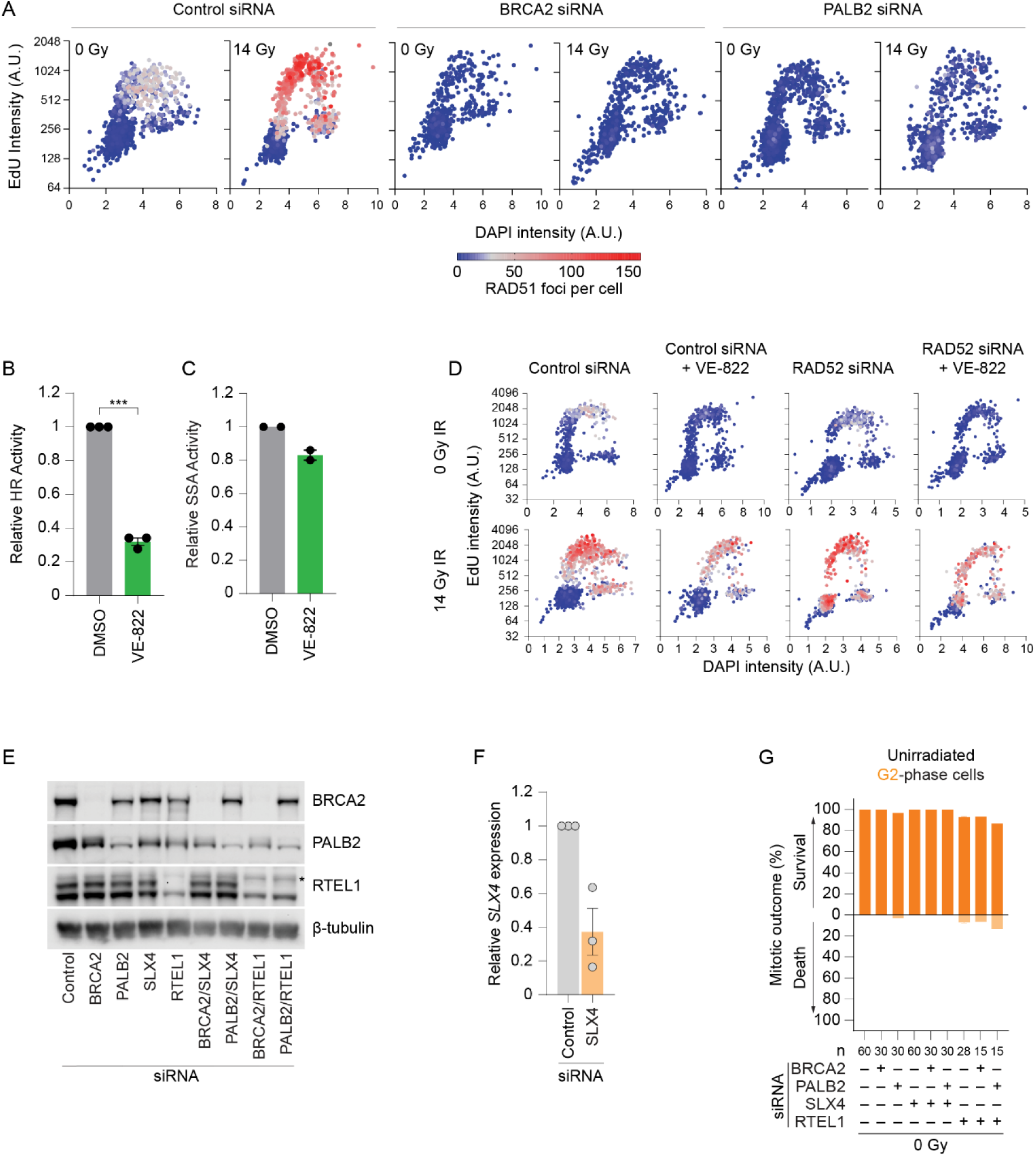
BRCA2, PALB2, and ATR promote immediate mitotic death. **A)** RAD51 foci in HeLa cells transfected with the indicated siRNAs ± 14 Gy IR imaged and presented as in Fig. 2F. **B, C)** HR (B) or SSA (C) repair of an I-SceI DSB repair reporter in HeLa cells ± 0.2µM VE-822 as measured by flow cytometry (mean ± SEM; n ≥ 2 biological replicates; paired t-test). Samples were collected 48 hours after I-SceI transfection and 40 hours after VE-822 addition. **D)** RAD51 foci for data shown in Fig. 3E. A representative biological replicate is shown. **E)** Western blot of whole cell extracts derived from 3F HeLa cultures transfected with the indicate siRNAs. **F)** RT-qPCR validation of SLX4 knockdown (n = 3 biological replicates). **G)** First mitosis outcome measured by live imaging following mock irradiation of synchronized G2-irradiated 3F HeLa cells transfected with the indicated siRNAs (mean ± SEM of 2 biological replicates; Fisher’s exact test). Where appropriate, *** p <0.001.

**Extended Data Figure 4:**
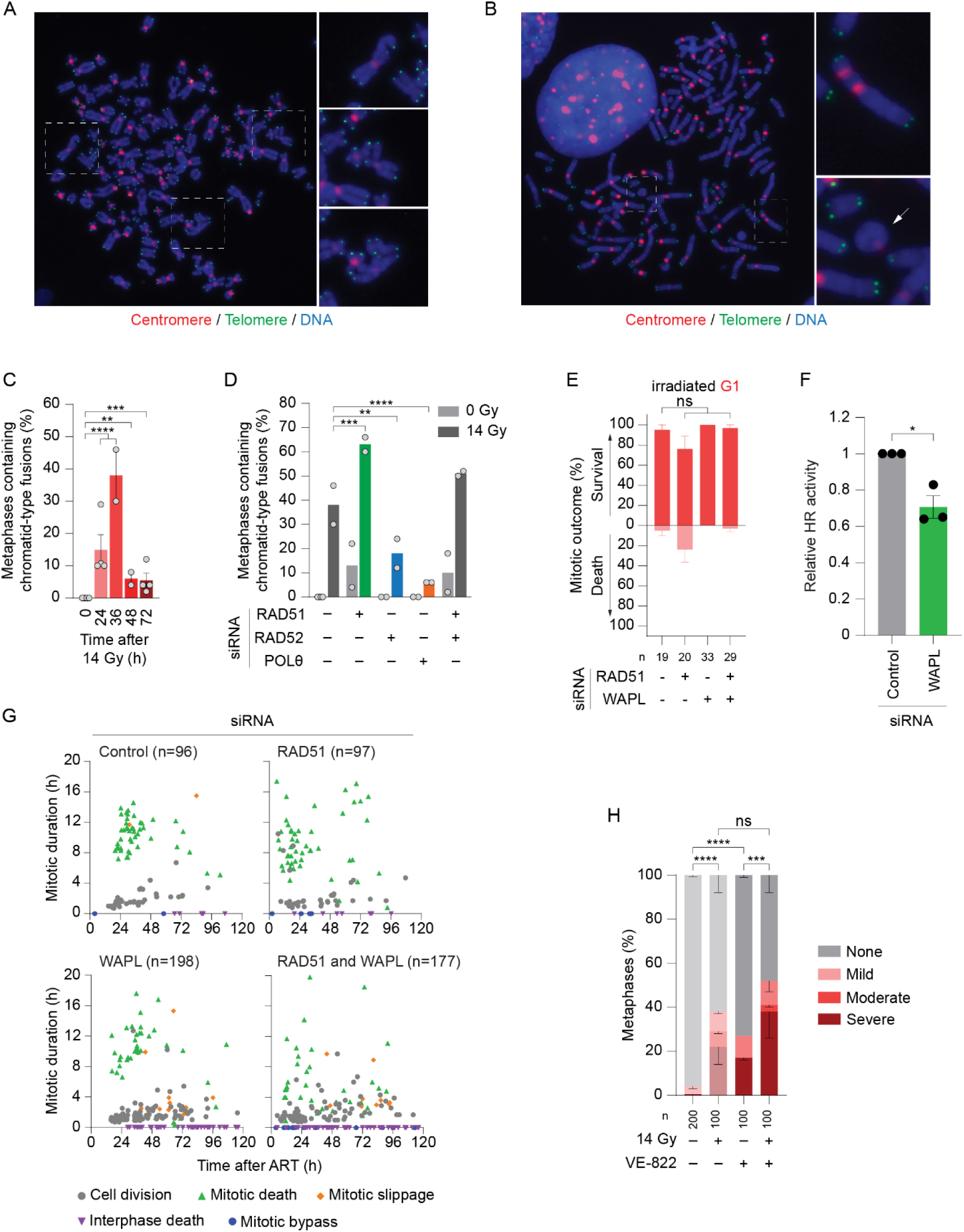
RAD51 and WAPL depletion differently rescue immediate mitotic death. **A, B)** Representative images of cytogenetic chromosome spreads depicting chromatid-type fusions (A), and ring and dicentric chromosomes (B) from HeLa cultures, prepared as in Fig. 4A. Expanded examples are shown on the right. **C)** Quantification of chromatid-type fusions depicted in (A) from control siRNA treated HeLa cells at the indicated times following 14 Gy IR (mean ± SEM; n ≥ 2 biological replicates scoring 50 metaphases per replicate; Fisher’s exact test). **D)** Quantification of chromatid-type fusions in cultures treated with the indicated siRNAs ± 14 Gy IR (mean ± SEM; n = 2 biological replicates scoring 100 metaphases per condition). The control siRNA data are the same data as the 0- and 36-hour time points in (C). **E)** Outcome of the first mitosis in 3F HeLa G1 phase cells irradiated with 14 Gy as determined by live imaging (mean ± SEM of ≥ 2 biological replicates; Fisher’s exact test). **F)** HR repair of an I-SceI DSB repair reporter in HeLa cells measured by flow cytometry (mean ± SEM; n = 3 biological replicates; paired t-test). **G)** Multi-dimensional representation of cellular outcomes in siRNA transfected 3F HeLa cultures following 14 Gy IR. Data from ≥ 2 biological replicates are pooled and presented as described in Fig. 1F. **H)** Quantitation of cohesion fatigue in HeLa cultures following mock or 14 Gy IR ± 0.2µM VE-822 (mean ± SEM of ≥ 2 biological replicates scoring 50 metaphases per replicate/condition; Fisher’s exact test). VE-822 was added immediately prior to mock or 14 Gy IR treatment. The VE-822 (-) samples are the same as in Fig. 4F. Where appropriate, ns = not significant, *p < 0.05, **p < 0.01, *** p <0.001, ****p < 0.0001. For panels E, G and H, n = the number of cells scored across all biological replicates.

**Extended Data Figure 5:**
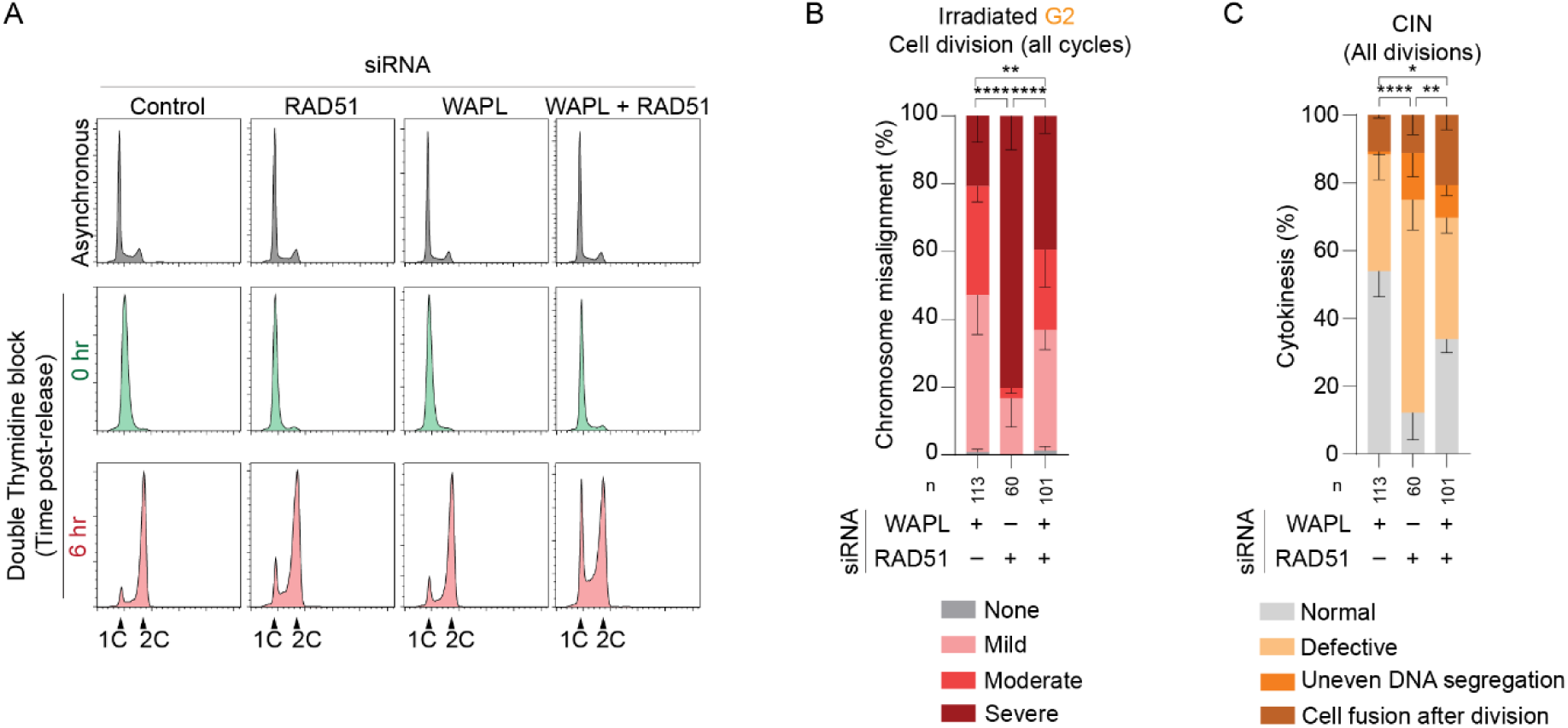
WAPL and RAD51 knockdowns have different consequences on genomic instability in cells refractory to mitotic death. **A)** Cell cycle profiles determined by flow cytometry in asynchronous, G1/S synchronized, and G2-enriched HeLa cultures following siRNA transfection. Cultures were G1/S synchronized using a double thymidine block. G2 enrichment is six hours after release from G1/S synchrony. Representative examples of ≥ 20,000 cells per condition are shown. **B)** Mitotic chromosome alignment from all mitoses that occurred within 120 hours after 14 Gy IR in G2-enriched mCherry-H2B HeLa cells as determined by spinning-disk confocal live imaging (mean ± SEM of 3 biological replicates; Fisher’s exact test). **C)** Chromosomal instability (CIN) and cytokinesis defects from cells that completed mitosis in (B) (mean ± SEM of 3 biological replicates; Fisher’s exact test). Where appropriate, *p < 0.05, **p < 0.01, ****p < 0.0001. For panels B and C, n = the number of cells scored across all biological replicates.

**Extended Data Figure 6:**
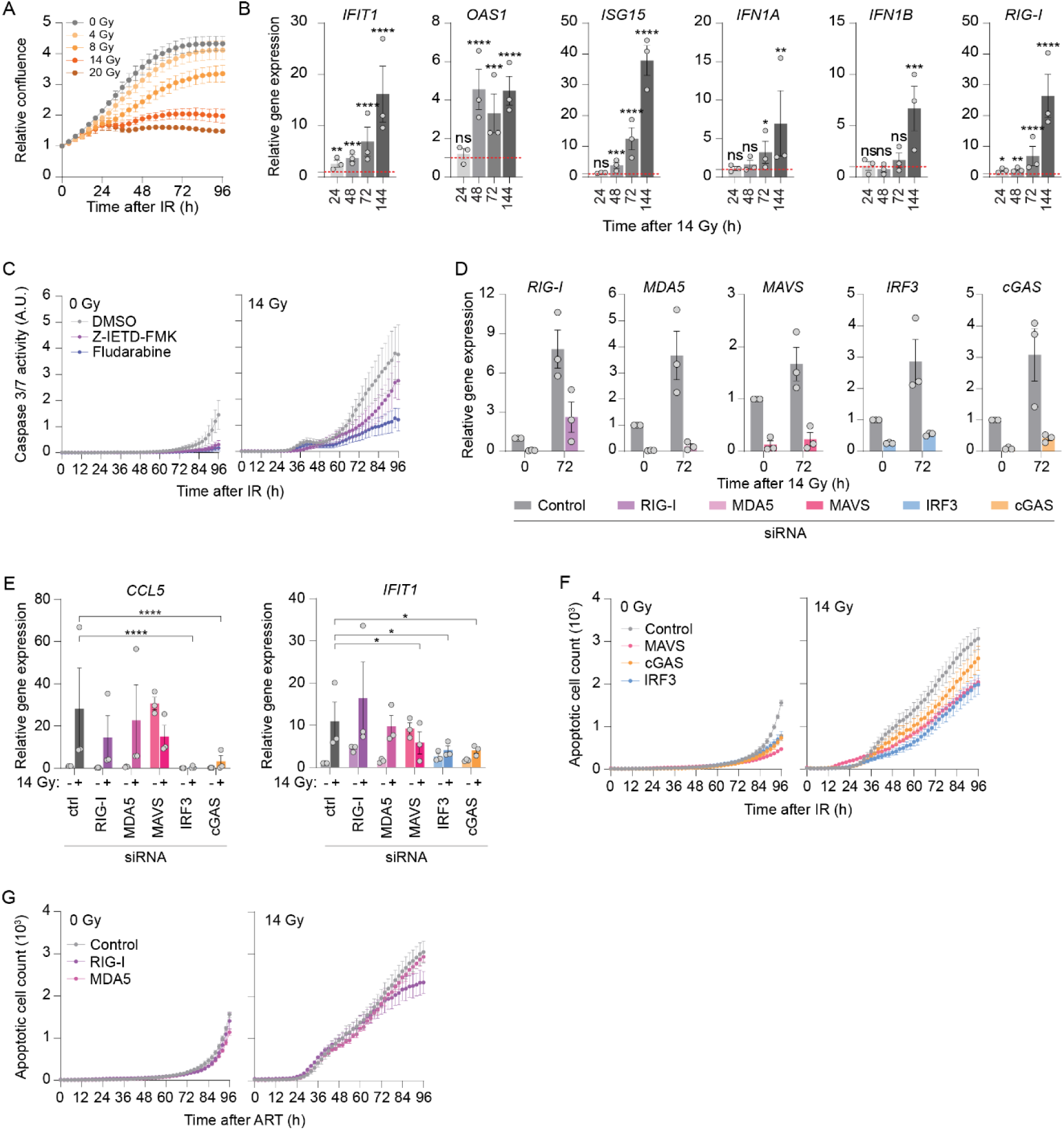
Distinct apoptotic pathways underpin cell cycle-regulated lethality during mitotic catastrophe. **A)** Growth curves of HeLa cultures treated with different IR dosages as measured through IncuCyte live imaging (mean ± SEM; n = 3 biological replicates). **B)** RT-qPCR measurement of ISG transcripts from HeLa cells at the indicated times following 14 Gy IR (mean ± SEM; n = 3 biological replicates; RM one-way ANOVA with Fisher’s LSD test). Fold change in gene expression is presented as relative to cells collected prior to IR (T0). The basal gene expression for non-irradiated cells is normalized to 1 and depicted as a red dotted line. **C)** Apoptosis measured through IncuCyte live imaging of a fluorescent cleaved Caspase 3/7 reporter in mock and 14 Gy irradiated HeLa cultures ± DMSO, 40µM Z-IETD-FMK or 10µM Fludarabine (mean ± SEM; n = 4 biological replicates). **D)** siRNA knockdown validation as measured through RT-qPCR (mean ± SEM; n = 3 biological replicates). Fold change in gene expression is relative to control siRNA treated cells collected immediately prior to IR (T0), normalized to 1. **E)** RT-qPCR measurement of *CCL5* and *IFIT1* expression in HeLa cultures following siRNA ± 14 Gy IR, presented as in (B) (mean ± SEM; n = 3 biological replicates, RM one-way ANOVA with Fisher’s LSD test; depicted p values are relative to ctrl siRNA + 14 Gy IR). **F, G)** Apoptosis measured as in (C) in siRNA transfected HeLa cultures ± 14 Gy IR (mean ± SEM; n ≥ 3 biological replicates). Where appropriate, ns= not significant, *p < 0.05, **p < 0.01, ***p < 0.001, ****p < 0.0001.

**Extended Data Figure 7:**
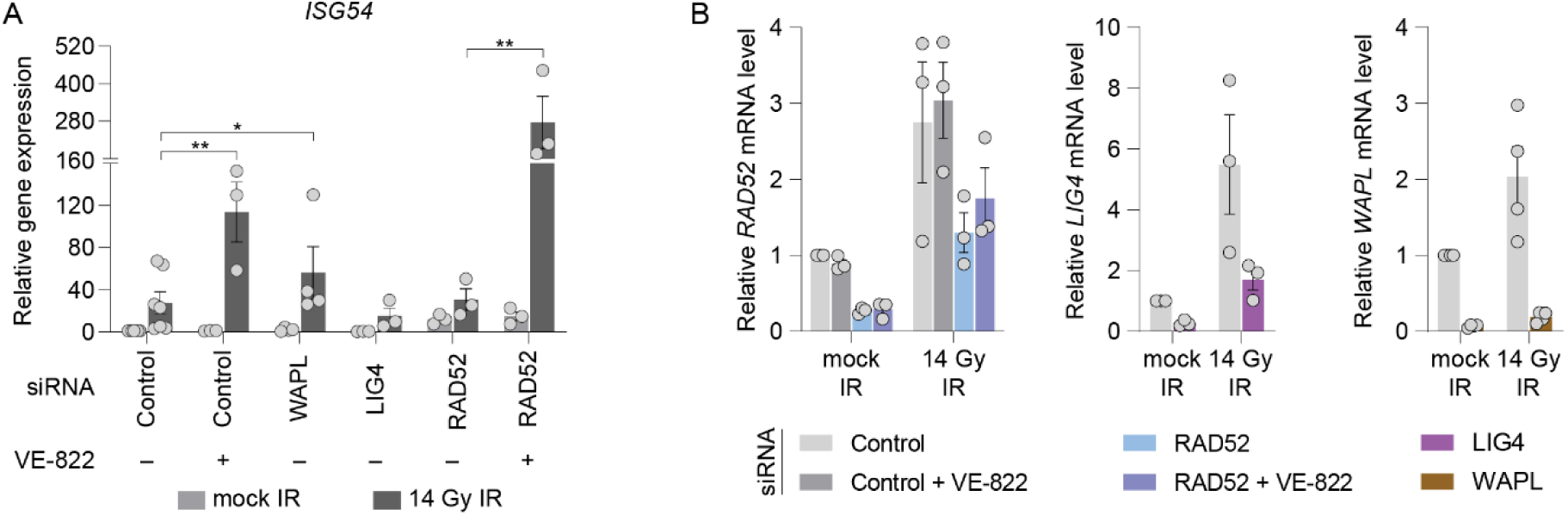
Pharmacological or genetic alteration of DSB repair impacts the innate immune response during mitotic catastrophe. **A)** RT-qPCR measurement of *ISG54* expression in HeLa cultures treated with the indicated siRNA ± 14 Gy IR and with or without 0.2µM VE-822 (mean ± SEM; n ≥ 3 biological replicates, Mixed-effect one-way ANOVA with Fisher’s LSD test). VE-822 was added an hour prior to mock or 14 Gy IR treatment. **B)** RT-qPCR validation of siRNA knockdown for the experiment in (A) and Fig. 7A (n ≥ 3 biological replicates). Where appropriate, *p < 0.05, **p < 0.01.

## SUPPLEMENTAL VIDEO LEGENDS

**Supplementary Video 1: Visualization of mitotic catastrophe.** Combined differential interference contract (DIC) and fluorescence live imaging of 3-colour FUCCI (3F) ^29^ HeLa cells treated with mock or 14 Gy IR. Cell cycle coloration as shown in Fig 1A. Time is relative to imaging commencement within one hour of mock or IR treatment.

**Supplementary Video 2: Cellular outcomes during mitotic catastrophe.** Examples of cellular outcomes observed with DIC live imaging in 3F HeLa cultures treated with mock or 14 Gy IR. Time is hrs:min relative to the start of the first image in the depicted outcome. Scale bars = 20µm.

**Supplementary Video 3: DNA-PKcs inhibition promotes mitotic death in G1 irradiated cells.** DIC and fluorescence live imaging of 3F HeLa cells treated with 20 Gy IR ± 0.5µM NU7441. Time is relative to imaging commencement within one hour of mock or IR treatment. NU7441 was added 30 minutes prior to irradiation.

**Supplementary Video 4: RAD51 depletion rescues immediate mitotic death of G2 irradiated cells.** DIC and fluorescence live imaging of G2-enriched 3F HeLa cells treated with control or RAD51 siRNA and 14 Gy IR. Cells were synchronized at G1/S boundary by double thymidine block, released for 6 hours, and irradiated in G2 phase as confirmed by 3F coloration. Time is relative to imaging commencement within one hour of mock or IR treatment.

**Supplementary Video 5: Chromosome segregation errors in cells that complete the first mitosis after genomic damage.** Spinning disc confocal live imaging of H2B-eGFP HeLa cultures treated with mock or 20 Gy IR. Time is shown as hrs:min relative to imaging commencement within one hour of mock or IR treatment. Scale bars = 20µm.

**Supplementary Video 6: WAPL and RAD51 depletion rescue immediate mitotic death during mitotic catastrophe through different mechanisms.** Combined spinning disc confocal and DIC live imaging of G2-enriched mCherry-H2B HeLa cultures treated with the indicated siRNA and 14 Gy. Cells were synchronized at G1/S boundary using a double thymidine block, released, and 14 Gy irradiated 6 hours later when the cells were in G2 (Fig. S5A). Time is relative to imaging commencement within one hour of mock or IR treatment. Scale bars = 20µm.

**Supplementary Figure 1: Flow cytometry gating strategies.** A-D) Homologous recombination (HR) reporter assays. For experiments using the pDRGFP reporter assay, outcomes were determined by gating A) live cell events followed by B) single cell events. Events were then gated for C) transfection efficiency using a generic mCherry expressing plasmid, or D) identified as GFP positive following transfection of an I-SceI expressing plasmid. **E-H)** Single strand annealing (SSA) reporter assays. For experiments using the hprtSAGFP reporter assay, outcomes were determined by gating E) live cell events followed by F) single cell events. Events were then gated for G) transfection efficiency using a generic mCherry expressing plasmid, or H) identified as GFP positive following transfection of an I-SceI expressing plasmid. I, J) Cell cycle analysis was performed by gating I) single cell events before J) plotting outcomes in a histogram. For all experiments >20,000 events were collected for each sample measured.

