## Supplementary Information for "Homologous recombination promotes mitotic death to suppress the innate immune response"

Supplementary Figure 1

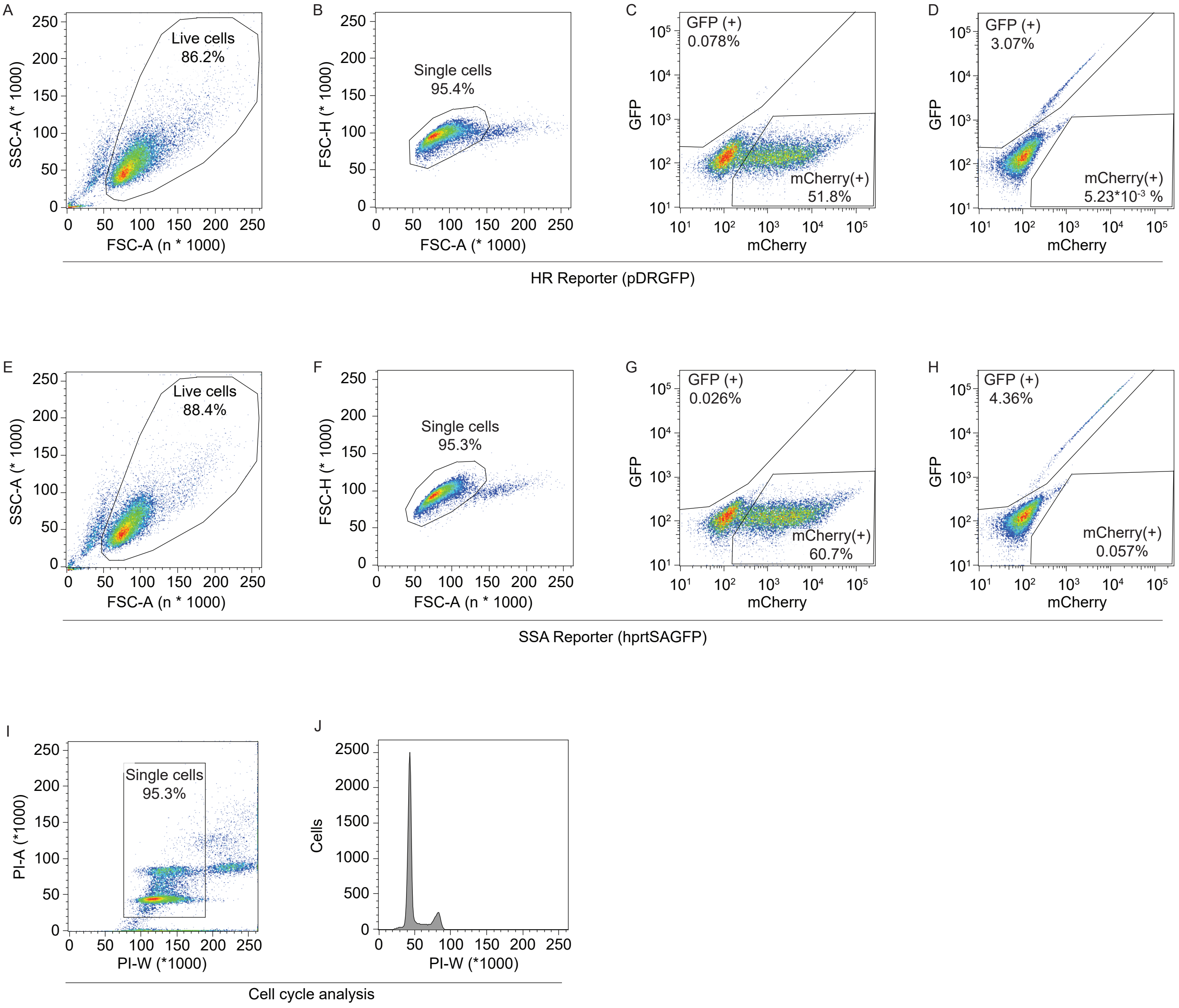

**Supplementary Table 1. Primers for molecular cloning**

| PRIMER | SEQUENCE (5' – 3') | SOURCE |
|---|---|---|
| H2B Forward | GATCCACCGGTCGCCACCATGCCAGAGCCAGCGAAG | This paper |
| H2B Reverse | ATTCCACAGCCGGATCTAAGCGCTGGTGTACTTGG | This paper |
| mCherry Forward | TCTAGGCGCCGGAATGGTGAGCAAGGGCGAGGAGG | This paper |
| mCherry Reverse | GGTGGCGACCGGTGGATCCTTGTACAGCTCGTCCATGC | This paper |
| pLXSN Forward | ATCCGGCTGTGGAATGTGTG | This paper |
| pLXSN Reverse | ATTCCGGCGCCTAGAGAAGG | This paper |
| crRNA sequence | GCCGATGGTGAAGTGGTAAG | Reference 36 |
| LBR2 Forward | AAATGGCTGTCTTTCCCAGTAA | This paper |
| LBR2 Reverse | GTAGCCTTTCTGGCCCTAAAAT | Reference 36 |

**Supplementary Table 2. Primers for RT-qPCR**

| PRIMER | SEQUENCE (5' – 3') | SOURCE | IDENTIFIER |
|---|---|---|---|
| CCL5 Forward | ACAGCCTCTCCCACAGGTA | This paper | N/A |
| CCL5 Reverse | TGTGGTGTCCGAGGAATATGG | | |
| cGAS Forward | AAGCAACTACGACTAAAGCCAT | | |
| cGAS Reverse | GATAGCCGCCATGTTTCTTCTTG | | |
| GAPDH Forward | AAGGTCGGAGTCAACGGATTTG | | |
| GAPDH Reverse | TGAGGTCAATGAAGGGGTCAT | | |
| IFIT1 Forward | CAGAACGGCTGCCTAATTTACA | | |
| IFIT1 Reverse | TCCCACACTGTATTTGGTGTCT | | |
| IFN1A Forward | GGAGGTTGTCAGAGCAGAAATC | | |
| IFN1A Reverse | ATAGCAGGGGTGAGAGTCTTTG | | |
| IFN1B Forward | TATGGGAGGATTCTGCATTACC | | |
| IFN1B Reverse | GGCTAGGAGATCTTCAGTTTCG | | |
| IRF3 Forward | CTCGTGATGGTCAAGGTTGTG | | |
| IRF3 Reverse | AATGTGCAGGTCCACAGTATTC | | |
| ISG15 Forward | TCTTTGCCAGTACAGGAGCTT | | |
| ISG15 Reverse | CAGGGACACCTGGAATTCGTT | | |
| ISG54 Forward | GAAGATTTCTGAAGAGTGCAGC | | |
| ISG54 Reverse | ATCAAGTTCCAGGTGAAATGGC | | |
| LIG4 Forward | CTGGAACTGTATTGCCTGCTTT | | |
| LIG4 Reverse | TGATGAATCTTCTCGTTTAACTGG | | |
| MAVS Forward | AGAGAGAAGGAGCCAAGTTACC | | |
| MAVS Reverse | AGGGCTTGCTCTGAATTCTCT | | |
| MDA5 Forward | ATGCAACCAGAGAAGATCCATT | | |
| MDA5 Reverse | AAACACGTTCTTTGCGATTTCC | | |
| OAS1 Forward | CTGGATTCTGCTGGCTGAAAG | | |
| OAS1 Reverse | TGTGCTGGGTCTATGAGAGAAA | | |
| POLQ Forward | CTGCTGTTTATGCAGGGATGAT | | |
| POLQ Reverse | CACAGTATGAAAGCCAGAAGCA | | |
| RAD51 Forward | ATTTCACGGTTAGAGCAGTGTG | | |
| RAD51 Reverse | GCTTTGGCTTCACTAATTCCCT | | |
| RAD52 Forward | GCTCAGTGTTATGCTTTGGACA | | |
| RAD52 Reverse | CCTCAATGTAGCACACCTTCTG | | |
| WAPL Forward | GATTCCCAGCACCATCAGAATC | | |
| WAPL Reverse | GAAGCATCTTGTTCCAGTTCCA | | |
| RIG-I Forward | TGCGAATCAGATCCCAGTGTA | Reference 92 | N/A |
| RIG-I Reverse | TGCCTGTAACTCTATACCCATGT | | |

**Supplementary Table 3. siRNAs used in this study**

| siRNA | SOURCE | IDENTIFIER |
|---|---|---|
| ON-TARGETplus Non-targeting Control SMARTpool | Dharmacon | Cat# D-001810-10 |
| ON-TARGETplus Human DDX58 (RIG-I) SMARTpool | Dharmacon | Cat# L-012511-00 |
| ON-TARGETplus Human IFIH1 (MDA5) SMARTpool | Dharmacon | Cat# L-013041-00 |
| ON-TARGETplus Human MAVS SMARTpool | Dharmacon | Cat# L-024237-00 |
| ON-TARGETplus Human IRF3 SMARTpool | Dharmacon | Cat# L-006875-00 |
| ON-TARGETplus Human MB21D1 (cGAS) SMARTpool | Dharmacon | Cat# L-015607-02 |
| ON-TARGETplus Human PRKDC (DNA-PKcs) SMARTpool | Dharmacon | Cat# L-005030-00 |
| ON-TARGETplus Human LIG4 SMARTpool | Dharmacon | Cat# L-004254-00 |
| ON-TARGETplus Human POLQ SMARTpool | Dharmacon | Cat# L-015180-01 |
| ON-TARGETplus Human BRCA2 SMARTpool | Dharmacon | Cat# L-003462-00 |
| ON-TARGETplus Human PALB2 SMARTpool | Dharmacon | Cat# L-012928-01 |
| ON-TARGETplus Human WAPL SMARTpool | Dharmacon | Cat# L-026287-01 |
| Silencer Select Negative Control #1 | Dharmacon | Cat# 4390843 |
| Silencer Select Human RAD51 | Ambion; Thermo Fisher Scientific | Cat# 4392420, s11735 |
| Silencer Select Human RAD52 | Ambion; Thermo Fisher Scientific | Cat# 4392420, s11747 |
